# Centromeric repeat diversity underlies non-Mendelian segregation pattern in hop (*Humulus lupulus*)

**DOI:** 10.1101/2024.11.03.621702

**Authors:** Lucie Horáková, Radim Čegan, Pavel Jedlička, Pavla Navrátilová, Hiroyuki Tanaka, Atsushi Toyoda, Takehiko Itoh, Takashi Akagi, Eiichiro Ono, Vojtěch Hudzieczek, Josef Patzak, Jan Šafář, Roman Hobza, Václav Bačovský

## Abstract

Aberrant meiosis in plants often leads to aneuploidy, genetic instability, and sterility. This can occur due to several factors, including chromosome misalignment, defective synapsis or environmental factors that may result in unusual genetic combinations in the offsprings. Unusual chromosome behavior during male meiosis in *Humulus lupulus* is linked to irregular chromosome segregation and genome instability. However, the origin of meiotic instability remains unclear.

We analyzed the centromeric landscape of *H*. *lup*ulus to determine its role in aberrant chromosomal segregation during cell division. Using a combination of bioinformatic, molecular and cytogenetic approaches, we identified new centromeric repeats and revealed two types of centromeric organizations. Cytogenetic localization on metaphase chromosomes confirmed the genomic distribution of major repeat arrays and revealed unique features that contribute to aberrant segregation.

Two centromeric types are composed of the major repeats SaazCEN and SaazCRM1 which are further accompanied by chromosome-specific centromeric satellites, Saaz40, Saaz293, Saaz85, and HuluTR120. Chromosome 2 displays unbalanced segregation during the cell division, implicating an important role for its centromere structure in segregation patterns. Moreover, Saaz293 is a new marker for studying aneuploidy in hop.

Our findings provide new insights on chromosome segregation in hop and highlight the diversity and complexity of the centromere organization in *H*. *lupulus*.

## Introduction

Centromeres are the sites of kinetochore assembly ensuring faithful segregation of chromosomes to daughter cells during mitosis and meiosis. Active regional centromeres are defined by the presence of the centromere-specific histone variant H3 (termed as CENH3 in plants). The CENH3-positive chromatin domain is localized within the primary constriction of monocentric chromosomes, allowing precise centromere localization in association with centromeric DNA and its functional relationship (Houben *et al*., 2007; Mendiburo *et al*., 2011; Maheshwari *et al*., 2017; Gent *et al*., 2017). Despite the evolutionary conservation of centromere function across eukaryotes (Henikoff *et al*., 2001), the amino acid sequence of CENH3, centromere size, and centromeric repeat sequences exhibit remarkable divergence among closely related species (Melters *et al*., 2013; Maheshwari *et al*., 2017). In plants, centromeric DNA is usually composed of a one or a few tandem repeats, organized into higher-order repeat structures, align with centrophilic long terminal repeat (LTR) retrotransposons, mostly belonging to Ty3/*Gypsy* superfamily (reviewed in Naish & Henderson, 2024). Retrotransposon-based centromeres have been identified in only few species, including einkorn wheat (Ahmed *et al*., 2023), apple (Zhang *et al*., 2019), and the moss *Physcomitrella patens* (Bi *et al*., 2024), highlighting rapid centromere evolution and diversity across plant kingdom (Lee *et al*., 2005). Centromeres are further embedded in pericentromeric chromatin, typically defined by histone modifications such as H3S10ph or H2At120ph, which contributes to the formation of a repressive chromatin state, having a crucial role in the kinetochore assembly (Neumann *et al*., 2016).

The role of (peri)centromere chromatin and kinetochore complex in atypical chromosome behavior has been described in interspecific plant hybrids of *H*. *vulgare* x *H*. *bulbosum* (Sanei *et al*., 2011) and *Festuca* x *Lolium* (Majka *et al*., 2023). In the latter study, both parental sexes were shown to influence non-Mendelian inheritance, illustrating the role of centromere structure and function in chromosome elimination and genome dominance. In addition to the essential centromere function, centromeres can experience a phenomenon known as centromere shift, where the original centromere relocates to a new position. Recent studies on einkorn wheat have shown that its centromeres are composed almost entirely of two major retrotransposons, *RLG_Cereba* and *RLG_Quinta*, which exhibit recent insertions in functional centromeres (Ahmed *et al*., 2023; Heuberger *et al*., 2024). As a result, structural chromosomal rearrangements have been found in at least four chromosomes in the einkorn genome have been found to undergo structural chromosomal rearrangements, where existing centromere have shifted to a new position (Ahmed *et al*., 2023; Heuberger *et al*., 2024). Beyond the centromere shift, another important evolutionary mechanism is centromere repositioning which involves the formation of *de novo* centromeres in different positions on the same chromosomes, accompanied by the inactivation of the original centromere. This phenomenon has been identified in various plant species, such as maize (Liu *et al*., 2015), *Arabideae* (Mandáková *et al*., 2020), and wheat (Zhao *et al*., 2023), among others. On a broader scale, centromere repositioning has been shown to modify recombination frequency, and gene expression driving karyotype diversity in plants (Mandáková *et al*., 2020).

*Humulus lupus* L. (common hop) is a perennial dioecious plant with heteromorphic sex chromosomes (20, XX in females and 20, XY in males; Winge, 1923). Its genome is relatively large (2.8 Gb), with a considerable amount of repetitive DNA (64%) and a high level of heterozygosity (Natsume *et al*., 2015; Padgitt-Cobb *et al*., 2023). Female inflorescences, known as hop cones, contain important secondary metabolites (bitter acids, polyphenols and terpenes) with antimicrobial properties, making hop a necessary commodity used in the brewing and pharmaceutical industries (Neve, 1991; Zanoli & Zavatti, 2008). Genus *Humulus* includes European (*H*. *lupulus* var. *lupulus*), North American (*H*. *lupulus* var. *lupoloides*, *H*. *lupulus* var. *neomexicanus*, *H*. *lupulus* var. *pubescens*) and Japanese (*H*. *lupulus* var. *cordifolius*) varieties and cultivars (Small, 1978), making this genus a valuable model for studying sex chromosome evolution and intraspecies sex chromosome genome dynamics.

Previous genetic studies of *H*. *lupus* have shown that a significant proportion of molecular markers deviate from expected Mendelian segregation ratios (Seefelder *et al*., 2000; McAdam *et al*., 2013; Zhang *et al*., 2017). Cytogenetic evidence has revealed the presence of multivalent chromosomal complexes, and unusual chromosome morphology during prophase I, that lead to the formation of anaphase bridges during male hop meiosis (Sinotô, 1929; Ono, 1955; Neve, 1958; Haunold, 1974; Zhang *et al*., 2017; Easterling *et al*., 2018, 2020). These abnormalities were demonstrated further as non-Mendelian segregation patterns, increasing genetic diversity and genome size. These instabilities, on the other hand, pose significant challenges for the breeding new cultivars, compounded by the absence of accessible tools, that would make the identification of chromosome instabilities possible.

Despite the availability of genomic resources and cytogenetic fata on chromosomal irregularities, the mechanism underlying non-Mendelian segregation in *H*. *lupulus* is still under debate. Earlier studies hypothesized that this this phenomenon could be a conserved genomic feature of *H*. *lupulus*, a consequence of breeding, or associated with an unusual centromere structure, even involving the presence of a dicentric chromosome. Thus, understanding of centromere function in *H*. *lupulus*, will be crucial for the selection and identification of stable parental lines, enhancing hop breeding programs and marker assisted selection. Beyond its practical applications, the study of centromere structure in *H*. *lupulus* may provide insights into genomic changes during the evolution of the *Cannabaceae* family.

To clarify whether the centromere organization affects non-Mendelian segregation of chromosomes in *H*. *lupulus* var. *lupulus*, we developed *H*. *lupulus* CENH3 antibody and analyzed centromeric landscape of autosomes and sex chromosomes. Utilizing the ChIP-seq data on the female and male genomes of *H*. *lupulus* and detailed karyotypic analysis, we identified novel centromeric repeats within centromeric subdomains and estimated recent insertion of CRM belonging to Ty3/*Gypsy* that associated with CENH3 domains. Aside, we identified an autosomal pair that is involved in non-Mendelian segregation, the formation of unviable microspores, unreduced cells, micronuclei, and reduced fertility. We propose a possible mechanism leading to non-Mendelian segregation patterns and processes affecting the mis-segregating autosomes. These new findings contribute to a deeper understanding of the complexity of the hop genome and the behavior of chromosomes during cell division.

## Materials and Methods

### Plant material

Female *H*. *lupulus* var. *lupulus* cv. Saaz - Osvald’s clone 72 (2n = 20, XX), male named “Liběšice” (Lib male), and other three male accessions (15246, 15249, 15276; 2n = 20, XY) used in this study (Supporting Information Table S1) were provided by the Hop Research Institute Co. Ltd. in Žatec (Czech Republic). All plants were grown in the greenhouse under 16h daylight/ 8h dark conditions at the Department of Plant Developmental Genetics in Brno (Czech Republic).

### Generation of *H*. *lupulus* CENH3 antibody

Gene coding CENH3 protein was searched in HopBase (Hill *et al*., 2017) and NCBI database using tblastn algorithm. The resulting sequence was compared to already published CENH3 library (Fig. S1). HlCENH3 primers were designed in Geneious Prime (2023.1.1) and synthesized in GeneriBiotech (Table S2). HlCENH3 gene was amplified using Q5 High-fidelity DNA polymerase (M0491S; NEB), following manufacturer instructions. Resulting PCR products were purified using QIAquick PCR Purification Kit (28104; QIAGEN) and used for blunt cloning using CloneJET PCR Cloning Kit (K1231; ThermoFisher). Sequencing primers (pJET1.2 F/R) were used in all reactions. Samples were sequenced in Macrogen (Amsterdam, Netherlands). Multiple sequence alignment and sequence conservation was verified in Geneious Prime (Fig. S2). The peptide for antibody synthesis was selected based on hydrophilic profile determined by Kyte-Doolittle algorithm with linear weight variation model (Kyte & Doolittle, 1982). The histone core domains were determined in HistoneDB 2.0 (Draizen *et al*., 2016). The antibody against the CENH3 protein of *H*. *lupulus* was custom raised using the peptide N-SPATTPKKAARTK-C. Peptide synthesis, immunization of rabbit to produce anti-HlCENH3 antibody, and final peptide affinity purification of antisera were performed by Genescript (USA).

### Indirect immunostaining

Interphase nuclei of *H*. *lupulus* Saaz female were prepared as described in Houben *et al*. (2003) and Bačovský *et al*. (2019) with some modifications. After fixation of roots in 4% paraformaldehyde in 1x PBS, roots were incubated in 2% PVP-40 (polyvinylpyrrolidone) and 1% Triton X-100 diluted in 1x PBS for 15 min on ice (Lunerová & Vozárová, 2023). For immunostaining, slides were incubated overnight at 4°C with HlCENH3 antibody (diluted 1:1000). After washing of the primary antibody, anti-rabbit secondary antibody (FITC-conjugated, diluted 1:200) in 1% blocking solution was used for 60 min at 37°C. Slides were washed in 1x PBS, dehydrated in an ethanol series, and mounted in Vectashield Antifade Mounting Medium supplemented with 4’,6-diamidino-2-phenylindole (DAPI). Images were captured using Olympus AX70 epifluorescence microscope equipped with cooled cube camera and processed using Adobe Photoshop. Immunostaining was performed in two independent experiments.

### Chromatin immunoprecipitation with sequencing (ChIP-seq)

Nuclei for ChIP experiments were isolated from young leaves of Lib male *H*. *lupulus*. The ChIP-seq protocol with anti-HlCENH3 antibody was performed as described in Navrátilová *et al*., (2022) in two biological replicates. The sequencing of the libraries was conducted on Illumina NovaSeq instrument with 150 bp paired-end reads on NovaSeq S1 flowcell (Illumina Inc., San Diego, CA, USA) at the Centre of Plant Structural and Functional Genomics in Olomouc, Czech Republic.

### Analysis of ChIP-seq data

The reads of HlCENH3-ChIP and input control were quality checked and filtered as genomic DNA using FastQC and Trimmomatic 0.32 (Bolger *et al*., 2014). To identify centromeric candidates and evaluate the enrichment of repetitive sequences in sequencing data from HlCENH3-ChIP experiment, we have applied the approach based on RepeatExplorer2 (Novák *et al*., 2013, 2020) and ChIP-seq Mapper tool (Neumann *et al*., 2012) similarly to Navrátilová *et al*. (2022), with the repeat contig sequences of *H. lupulus* identified by RepeatExplorer2 as a reference. For the further analyses, we have used either consensus sequence of the candidate cluster (if available) or the individual contigs from the candidate cluster identified by blastn search (if the RepeatExplorer2 output did not show consensus sequence of the cluster). In parallel the trimmed reads of input and HlCENH3-ChIP were used for centromere identification and characterization in Saaz genome (accession number: AP031542-AP031551, AP035121-AP035448) and male cultivar “10-12” (accession number: AP031552-AP031562, AP031914-AP032066). Briefly, reads were mapped by bwamem2 (Vasimuddin *et al*., 2019) to female and male genome of *H. lupulus* and analyzed by MACS3 (Zhang *et al*., 2008). The most enriched regions on each chromosome were considered as the centre of the centromeric region and from this site we extracted 3 Mbp upstream and downstream (6 Mbp) for detailed analysis.

### Repeat annotation in centromeric sections of *H. lupulus* genome assembly

We used *H. lupulus* assembly sections for either mapping of RepeatExplorer generated repeat clusters using RepeatMasker (version 4.1.1; Tarailo-Graovac & Chen, 2009) for identification of intact LTR retrotransposons insertions using DANTE_LTR (https://github.com/kavonrtep/dante/). The insertion age was estimated using Long Tandem Repeats divergence (Jedlicka et al. 2020) and synonymous substitution rate 6.1×10^−9^ (Padgitt-Cobb *et al*., 2023). In order to distinguish possible lineages of autonomous and non-autonomous CRM Ty3/*Gypsy* elements, the all intact CMR transposons were aligned using MAFFT version 7 (Katoh & Standley, 2013) and Maximal-Likelyhood tree was generated using FastTree 2 (Price *et al*., 2010).

### Preparation of DNA probes

Total genomic DNA of *H*. *lupulus* was isolated from young leaves using the NucleoSpin Plant II (740770-50; Macherey-Nagel GmbH and Co. KG.), according to the manufacturer’s instructions. The primers for centromeric repeats were designed using Geneious Prime (2023.1.1) based on the RepeatExplorer2 consensus sequences. The primer sequences are listed in (Table S2). DNA probes were amplified using PCR in a mixture containing 1x PCR buffer, 0.0001 M dNTPs, 0.0001 M of each primer, 0.5 U Taq polymerase (Top Bio), 10-15 ng of template DNA in a total of 20 µl. The PCR cycling conditions followed manufacturer instructions (95°C for 4 min followed by 35 cycles of 94°C for 30 s, 55-60°C for 35 s, 72°C for 30 s and final extension step of 72°C for 10 min). The annealing temperature was optimized for each primer pair. PCR products were separated on 1.5% agarose gel with EtBr staining and purified using the QIAquick PCR Purification Kit (28104; QIAGEN). Purified DNA (1 µg) was labeled with Atto488 NT (PP-305L-488), Atto550 NT (PP-305L-550) or Cy5 (PP-305L-647N) using Nick translation labeling kits (Jena Bioscience) following manufacturer instructions. After 90 min at 15°C, the NICK translation products were checked on 1% agarose gel with EtBr staining. The reaction of well-labeled DNA probes was stopped by addition of 0.5M EDTA and incubation on 85°C. The labelled probes were further used directly to hybridization mixture.

### Mitotic and meiotic chromosome preparation

Mitotic chromosomes were prepared as described in (Razumova *et al*., 2023) with some modifications. Young leaves (2-5 mm in length) were collected from intensively grown female and male plants of *H*. *lupulus*. The leaves were pretreated with 0.002M 8-hydroxyquinoline for 4 h (2 h at RT and 2 h at 4°C in the dark) and then fixed in Clarke’s fixative (ethanol:glacial acetic acid, 3:1, v/v) at 4°C overnight. The fixative was replaced with 70% ethanol and the leaves were used directly for squashing (or stored at −20°C until use). Fixed leaves were washed 1x 5 min in distilled water, 1x 5 min in 45% acetic acid, 2x 5 min in 0.001M citrate buffer and macerated in 1% enzyme mixture (Table S3) diluted in 0.001M citrate buffer for 50 min at 37°C. Leaves were squashed in 60% acetic acid. After freezing in liquid nitrogen, the coverslip was removed, and slides were incubated in freshly prepared Clarke’s fixative for 3-5 min. Prepared slides with well-preserved metaphase were used for FISH or stored at −20°C in 96% ethanol until use.

Male panicles were fixed in Clarke’s fixative for 24 h at room temperature, the fixative was replaced, and material was used for squashing method or stored at −20°C until used. Single anthers were washed 2x 5 min in 0.001M citrate buffer and macerated in 1% enzyme mixture (Table S3) diluted in 0.001M citrate buffer for 30 min at 37°C. Anthers were squashed in drop of 45% acetic acid. After freezing in liquid nitrogen, the coverslip was removed. Slides were incubated in freshly prepared Clarke’s fixative for 3-5 min. Meiotic chromosomes were counterstained by DAPI in Vectashield Antifade Mounting Medium.

### Fluorescence *in situ* hybridization (FISH)

FISH was performed on mitotic metaphase chromosomes as described in (Sacchi *et al*., 2024). The hybridization mixture (stringency 77%) contained 50% formamide, 10% dextran sulphate, 2x SCC and 1.5 ng/µl of each probe per slide. Chromosomes were counterstained by DAPI in Vectashield Antifade Mounting Medium. Images were captured using Olympus AX70 epifluorescence microscope equipped with CCD camera and processed using Adobe Photoshop. FISH with specific DNA probes was performed in at least three individual experiments and at least ten metaphases were analyzed per experiment.

### Pollen grain viability

Male panicles of three plants Lib male, F_1_ progeny (Osvald’s clone 72 x Lib male), and wild hop grown in Brno-Horní Heršpice (Table S1) were fixed in Carnoy’s fixative (ethanol:chloroform:acetic acid, 6:3:1, v/v). Pollen tetrads and pollen grain were stained according to (Peterson *et al*., 2010) with minor modifications. Fixed anthers were squeezed in the staining solution. Slides with anthers were heated on a hot plate at 83°C for 10 min. Images of pollen tetrads and pollen grains were captured using an Olympus CX43 microscope. The number of tetrads produced by microsporogenesis and the number of viable (purple) and non-viable (gray) pollen grains were counted in ImageJ FiJi (1.52) using Multi point Tool function.

## Results

### Analysis of centromeric landscape in *H*. *lupulus* and PAR position

To investigate the centromeric structure of female and male *H*. *lupulus*, we developed *H*. *lupulus*-specific antibody targeting the centromeric histone variant of histone H3, designated as HlCENH3 (Figs S1, S2). Immunostaining of HlCENH3 revealed twenty round-shaped like structures in the interphase nucleus, correlating with number of chromosomes observed in *H*. *lupulus* Saaz female and Lib male (2n = 20; Fig. S3). This supports previous observations of monocentric organization of *Humulus* chromosomes. The validated functional HlCENH3 antibody was used for ChIP-seq analysis. We found monocentric localization of HlCEHN3 enrichment in all ten chromosome pairs of *H*. *lupulus* (Figs 1, S4, S5). These regions and their surrounding areas resulted in 6 Mb-long regions, which we classified as centromeric and pericentromeric domains for further detailed analysis. ChIP-seq data analysis using RepeatExplorer2 (Novák *et al*., 2013, 2020) and ChIP-seq Mapper tool (Neumann *et al*., 2012), following previously established protocols, revealed six centromere-specific repeat candidates – SaazCEN, SaazCRM1, Saaz293, HuluTR120, Saaz85 and Saaz40 (Table S4). One of these repeats (Saaz293) was further supported by additional analysis of repetitive DNA composition in both male and female genomes of *H*. *lupulus* (Methods S1; Table S5-S6; Notes S1). The centromeric localization of these six newly identified repeats was further confirmed by FISH on metaphase chromosomes in both Saaz female and Lib male accessions (Fig. 1). Additionally, the previously published HSR1 subtelomeric repeat (Divashuk *et al*., 2011), which matches cl124 (Table S6; Notes S1), was used to distinguish the sex chromosomes of *H*. *lupulus*. The distribution of HSR1 supports previously described patterns in both sexes (Fig. 1b). Two newly identified repeats, designated in this study as SaazCEN and SaazCRM1, were enriched in centromeric region of all chromosomes, including X and Y chromosomes (Fig. 1b,c), except for chromosome 2 which showed lower densities for both repeats (Fig. S5). The position of centromeric-specific repeat SaazCEN suggests that *H*. *lupulus* karyotype consists of four metacentric chromosomes (1, 4, 7 and 9), five submetacentric chromosomes (2, 3, 5, 8, and X), and two acrocentric (6 and Y; Fig. S6). The centromere of chromosome 2 is primarily enriched for Saaz293 satellite (Figs 1a,d, S5). Additionally, the pericentromeric region of this chromosome is composed of Cl6 (Fig. 1e,f), a satellite with high sequence similarity to GenBank database accession MN537570, described as satellite HuluTR120 (Easterling *et al*., 2020). Physical localization of HuluTR120 revealed additional enrichment on chromosome 3, but only on one chromosome of the homologous pair (Fig. 1e,f). Comparison of three additional male accessions revealed differences in the distribution of HuluTR120 indicating intraspecies variability (Fig. S7). The Lib (Fig. 1e) and 15246 (Fig. S7a) male accessions display HuluTR120 in pericentromeric region similarly to Saaz female, only on one chromosome 3 of the homologous pair. In contrast, male accessions 15249 and 15276 (Fig. S7b,c) possess even number of HuluTR120 on both homologous chromosomes implicating segmental aneuploidy or hybrid constitution of chromosome 3. Furthermore, HuluTR120 shows strong enrichment along the whole (peri)centromeric region of the Y chromosome (Fig. 1e,f). This pattern is consistent in all three male accessions 15246, 15249, and 15276 (Fig. S7), confirming an ancestral origin of HuluTR120 satellite on the Y chromosome. Similar to chromosome 2, the Y chromosome showed lower densities of SaazCEN and SaazCRM repeats (Fig. S5). The centromere of chromosome 6 contains, in addition to SaazCEN also the satellites Saaz85 (Fig. 1g) and Saaz40 (Fig. S8a,b). FISH localization of Saaz40 showed similar results to Saaz85 even under lower stringency conditions (68%; Fig. S8a,b). This chromosome simultaneously possesses 45S rDNA on the p-arm in both sexes (Fig. S9a,b).

**Fig. 1.**
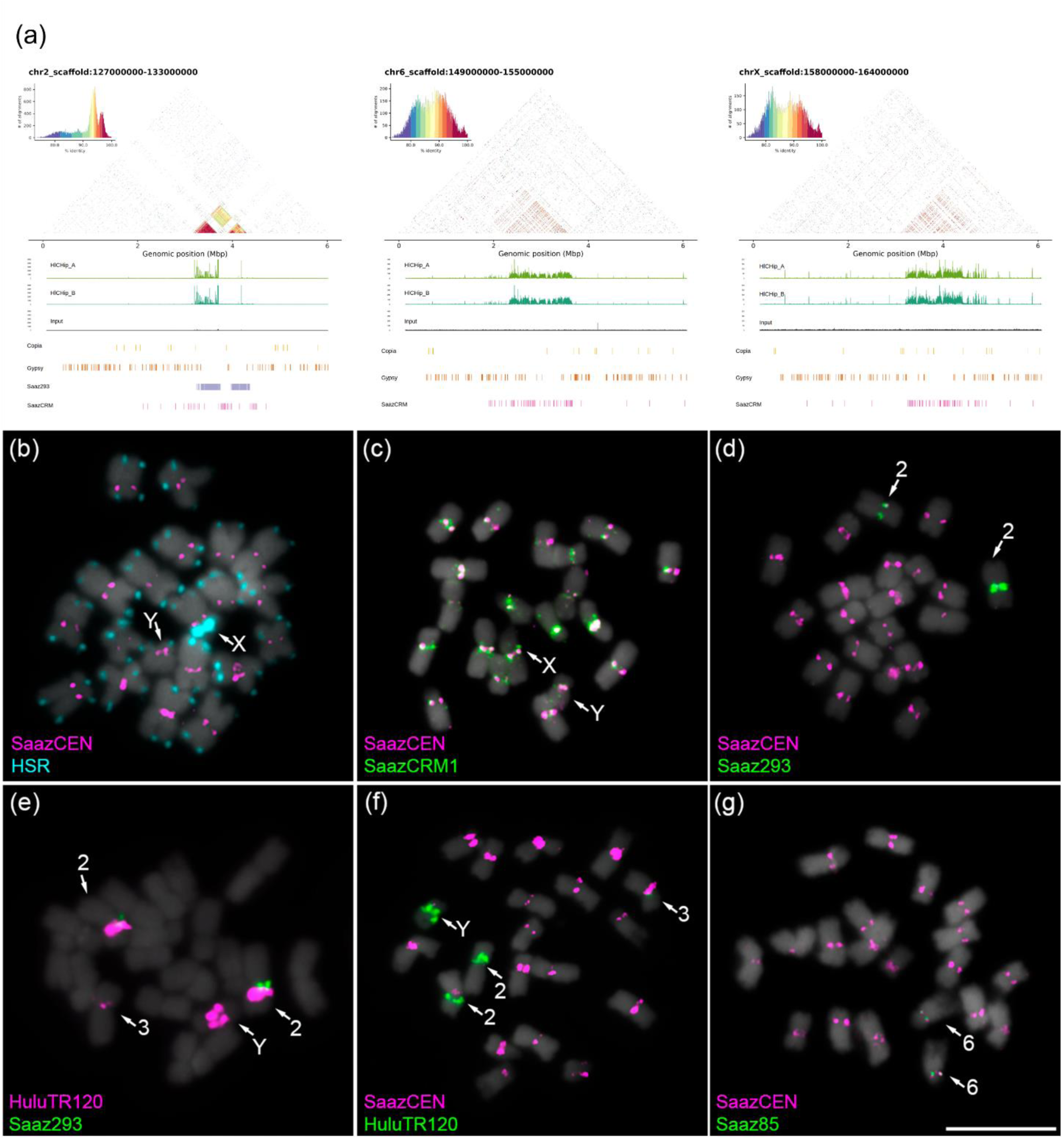
Characterization of the centromeric regions in *Humulus lupulus*. (a) Centromere characterization of chromosomes 2, 6, and X in *H*. *lupulus*. Heatmaps display pairwise sequence identity with dot plots revealing organization on all three chromosomes colored by percent identity. The first three lines compare HlCENH3 ChIP-seq data replicates (green and blue) against the input control. The subsequent lines show the detailed distribution of LTR-retrotransposons (Ty1/*Copia*, Ty3/*Gypsy*, and SaazCRM1) and chromosome-specific centromeric satellites Saaz293, Saaz40, Saaz85, and HuluTR120. Notably, chromosomes 2 and Y show a unique satellite array pattern. (b) Distribution of centromeric and centromere-related repeats on metaphase chromosomes of male *Humulus lupulus*. The centromeric repeat SaazCEN (magenta) together is shown with the subtelomeric tandem repeat HSR1 (cyan). The HSR1 repeat allows to distinguish X and Y chromosomes (arrows). HSR1 is present in the subtelomeric region of all autosomes, except the p-arm of chromosomes 6, and X chromosome (HSR1 signal in pericentromeric region). The Y chromosome displays HSR1 satellite in subtelomeric region of the p-arm. (c) Distribution of retrotransposon SaazCRM1 (green) and its colocalization with SaazCEN (magenta) in centromeric regions of all chromosomes. (d) Saaz293 satellite (green) colocalization with SaazCEN (magenta) in the centromeric region of chromosome 2. (e) Colocalization of Saaz293 satellite (green) and HuluTR120 satellite (magenta) on chromosome 2. (f) Localization of the HuluTR120 satellite (green) in the pericentromeric region of chromosome pair 2, chromosome 3, and Y. The Y chromosome displays two interstitial centromeric signals. (g) The Saaz85 satellite (green), together with the SaazCEN (magenta), constitutes the centromere of chromosome 6. Mitotic chromosomes were counterstained with DAPI. Scale bar = 10 µm.

The positions of SaazCEN and SaazCRM1 indicate that HSR1 locus (previously linked to the pseuodoautosomal region, PAR) is located on the short arm of the Y chromosome (Fig. 1b,c). This locates the PAR on the p-arm instead of on the q-arm. During diakinesis, the X and Y chromosomes are associated in an end-to-end bivalent conformation, with a subtelomeric HSR1 probe at the chromosomal ends (Fig. S10a) and HuluTR120 on the Y chromosome (Fig. S10b). This bivalent conformation further supports PAR localization on p-arm of the Y chromosome. No differences were observed in the distribution of all repeats on autosomes between sexes (Fig. S11). For clarification, all metaphase figures with HSR1 probe are shown in Fig. S12.

### Two centromere types in *H*. *lupulus* genome organization

The ideogram illustrates the overall distribution of *Humulus* repeats, including HSR1, 5S rDNA, 45S rDNA, and major centromeric repeats newly identified in this work, on the metaphase chromosome of Lib male of *H*. *lupulus* (Fig. 2a). The respective proportion of each centromeric repeats for each chromosome, including sex chromosome, defines two centromere types (Fig. 2b). The first type predominantly contains two major repeats - SaazCEN and SaazCRM1 (Fig. S5). The second type additionally includes Saaz293, Saaz85, and Saaz40 arrays, particularly on chromosome 2, 3, 6, 8 (Figs 2b, S5), and HuluTR120 present specifically on the Y chromosome (Figs 1e,f, 2b, S7, S10). However, physical localization defines Saaz293 as a centromere-specific satellite for chromosome 2 (Fig. 1d), and Saaz40 and Saaz85 as specific for chromosome 6 (Figs 1g, S8). A dot plot diagram of these three satellites revealed 50-60% sequence similarity (Fig. S13). The physical localization and sequence comparison indicated that HuluTR120 is in centromeric regions of chromosomes 2, 3, and Y (Figs 1e,f), further specifying HuluTR120 of chromosomes 2 and 3 respectively into pericentromeric region (Figs 2b, S5), whereas on the Y chromosome this satellite is highly enriched in the centromeric region (Fig. 2b). Compared to other autosomes or sex chromosomes (Figs 1a, S4), the strongest enrichment of Saaz293 is within the centromere of chromosome 2 (Figs 1a, 2b, S5). Based on the heatmap identity of the entire centromeric region (Fig. 1a), chromosome 2, 6 and Y display unique higher-order repeat satellite structure compared to the other autosomes, or X chromosome. This organization suggests crucial role of this satellite in centromere organization, and potential involvement in chromosome segregation.

**Fig. 2.**
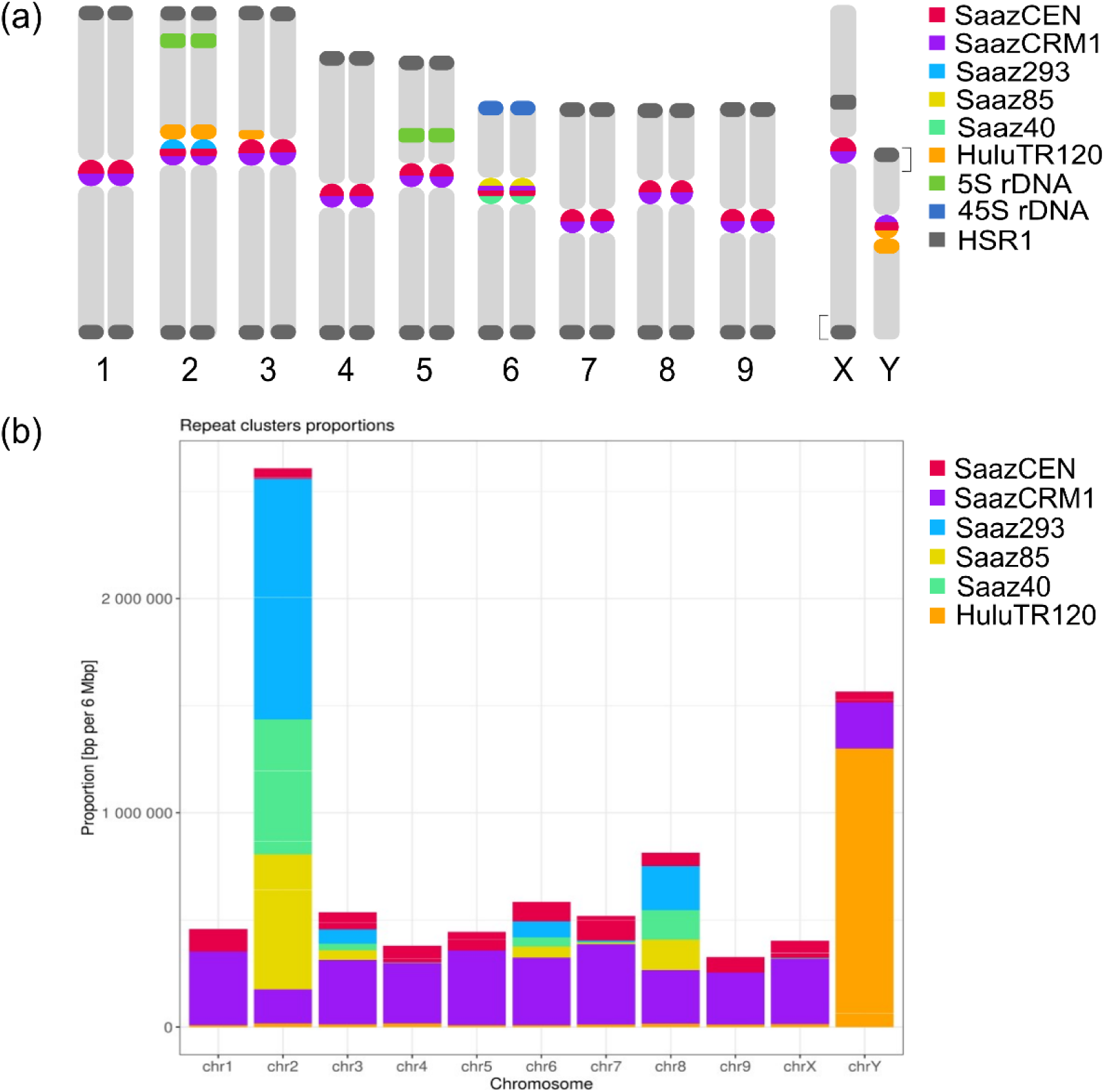
Proportion of identified tandem repeats and LTR retrotransposon within the centromeres of *Humulus lupulus*. (a) Ideogram of the *H*. *lupulus* showing FISH localization of centromeric repeats and other *Humulus*-specific repetitive sequences (Figs 1, S8-S12). Brackets indicate the positions of pseudoautosomal regions on the sex chromosomes. (b) Proportion of centromeric repeats (bp per 6 Mbp) for each of the autosome and sex chromosome. Centromeres of all chromosomes, including sex chromosome are composed of SaazCEN (red) and SaazCRM1 (purple) repeats. Notably, the centromere of chromosomes 2 and Y show enrichment for large satellite arrays compared to the other autosomes. This is particularly evident in the presence of Saaz293, Saaz40, and Saaz85 on chromosome 2, as well as HuluTR120 on the Y chromosome.

### Diversity of LTR retrotransposons in *Humulus* centromeres

The newly identified centromeric repeat SaazCEN is characterized by its insertion into LTR domains that are flanking central region of CRM retrotransposons. The SaazCEN basic monomer units is 284 bp, containing several 39 bp tandem repeats at the 3‘ terminus. The number of copies varies between SaazCEN arrays. We did not observe any consistent patterns among individual centromeres, regarding the number of copies for SaazCEN 39bp subunits.

From available sequence data, we found five clades (Ale, Angela, Ikeros, SIRE, and TAR) of the Ty1/*Copia* family and TEs from chromovirus (Tekay and CRM) and non-chromovirus (Retand and Athila) lineages of the Ty3/*Gypsy* family in all centromeres (Fig. S14a,b). Among Ty1/*Copia* we found Angela to be the most abundant clade, although Ty1/*Copia* exhibited overall lower copy numbers compared to Ty3/*Gypsy* families. The lowest proportion of Ty1/*Copia* family was found in the centromere of the Y chromosome (Fig. S14a). Within the Ty3/*Gypsy* family, Tekay and CRM, which possess chromodomains at the integrase C-terminal region (Neumann *et al*., 2019), were the most dominant sequences (Fig. S14b). Based on LTR similarity, the youngest CRM retrotransposon copies were found directly in HlCENH3 binding domains, with average insertion time being estimated between 0.0 to 1.0 Mya (Fig. S15). In contrast, Tekay elements are generally older with insertion ages between 0.0-10.0 Mya and are more dispersed within the centromeric region (Fig. S15). Interestingly, chromosomes 1, 2, 3, 5, 7, X, and Y display expanded HlCENH3 binding domain, based in ChIP-seq analysis (Figs S5, S15). The high density of recent CRM insertions in the immediate vicinity of HlCENH3 of chromosomes 1, 5 and Y, indicates an expansion of CRMs and potential shift in centromere position (Fig. S15).

### Autonomous and non-autonomous CRMs in *Humulus* centromere

We identified a total of 671 CRMs copies in *Humulus* centromeres, categorized into autonomous and non-autonomous CRM retrotransposons (noaCRM). Autonomous CRMs are composed of one large open reading frame (ORF) that encodes all canonical proteins required for retrotransposition. These include GAG protein, which forms virus-like particles in which reverse transcription takes place, reverse transcriptase (RT), RNAse H (RH), integrase (INT), chromodomain (CHDCR) and protease (Fig. 3a). The dominant noaCRM in *H*. *lupulus* based on the result in this work, lack essential proteins namely RT, RH, and INT (Fig. 3a; Langdon *et al*., 2000; Nagaki *et al*., 2005). Minor noaCRMs, a subclass of dominant noaCRM in *H*. *lupulus*, lack further protein resulting in incomplete ORF (Figs S16, S17). Autonomous and noaCRMs represent 189 and 482 of all identified CRMs, respectively (Fig. S17). Among noaCRMs, 446 copies are dominant, while 36 are minor elements.

**Fig. 3.**
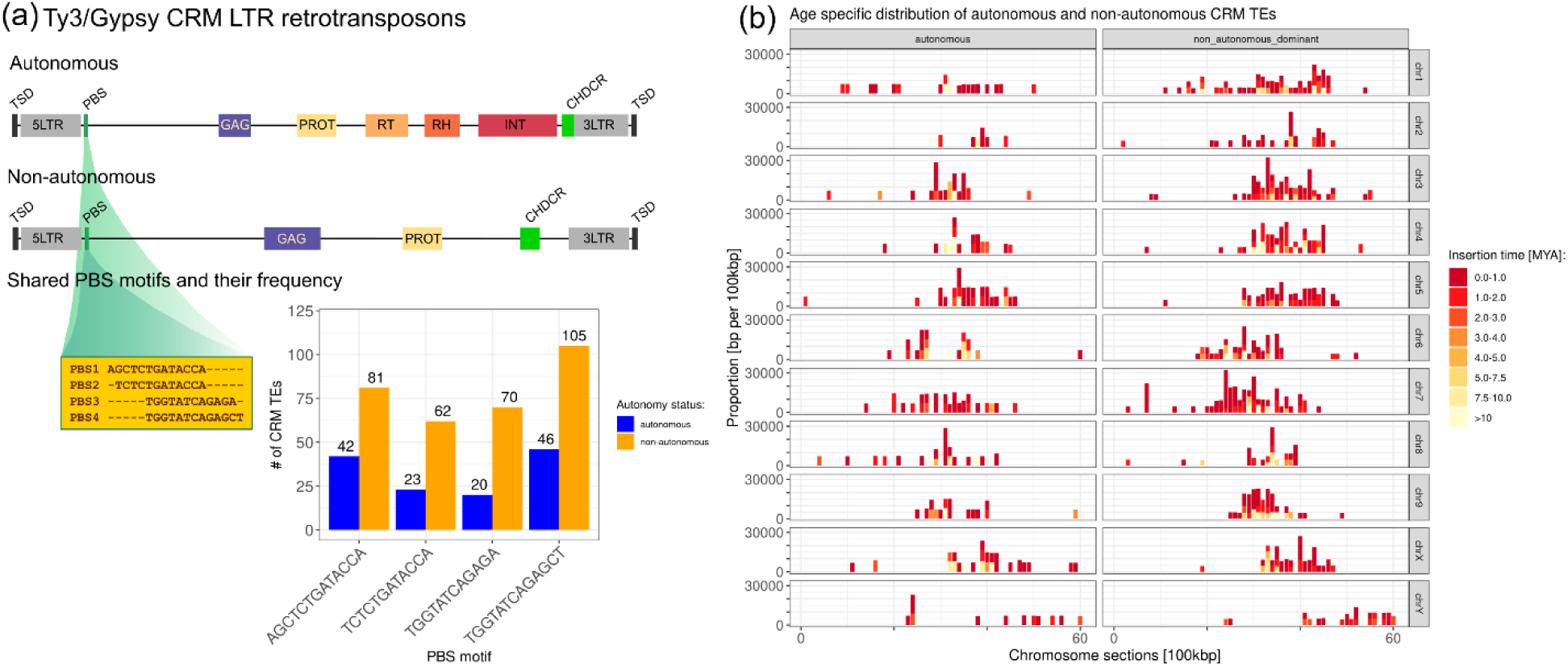
Characterization of CRM retrotransposons in the centromeres of *Humulus lupulus*. (a) The structures of autonomous and non-autonomous centromeric CRM retrotransposons in *Humulus lupulus*. Both retrotransposons have two direct long terminal repeats (5‘ LTR and 3‘ LTR) and primer binding site (PBS). Shared PBS motifs are shown in a yellow box. PBS4 is the most frequent in non-autonomous CRM. Autonomous CRMs contain six protein domains: GAG, protease (PRO), reverse transcriptase (RT), RNase H (RH), integrase (INT), and chromodomain (CHDCR type). In contrast, non-autonomous CRMs lack all three domains (RT, RH, and INT). Both retrotransposons are flanked by target site duplication (TSD). (b) Distribution of autonomous and non-autonomous (dominant) CRM retrotransposons and their insertion time (color scale from red to yellow) across *Humulus* centromeres. Note the recent insertions of both CRM groups in the positions of CENH3 enrichment.

From 671 CRMs, the primer binding site PBS motif was detected in 169 and 374 (i.e. 80.9%) of autonomous and noaCRMs, respectively. The sequence structure of the PBS and its comparison between noaCRMs and autonomous CRM showed that from 543 TEs, 449 (82.7%) belongs into one of the four dominant PBS motifs. Sequences of 5‘ LTR in groups with similar PBS motifs are identical in both autonomous and noaCRMs (Fig.3a), suggesting that reverse transcription and even replication may be triggered by the same mechanism in both CRMs groups. These findings provide further evidence of autonomous CRM-dependent retrotransposition of noaCRMs.

Comparison of insertion time and CRM distribution across individual centromeres did not reveal large differences between autonomous and non-autonomous categories (Fig. 3b). Phylogenetic and clustering analysis of autonomous CRMs and noaCRMs, combined with insertion time data, indicate that the most recent insertions (0.0-1.0 Mya) occurred within noaCRMs, or in subdomains of autonomous CRMs (Fig. S18a,b). Finally, SaazCEN repeats are present in the majority of both CRM categories (94.4%), including autonomous and noaCRMs (Fig. S19). Based on the HlCENH3 affinity and the localization of CENH3 binding domain within CRM, we found that HlCENH3 binding domains predominantly localize to non-coding regions of both CRM categories, with the second highest frequency of interactions occurring within LTR domains (Fig. S20).

### The effect of centromeric variability on chromosome segregation

The somatic aneuploidy or aneusomaty describe the nuclear condition of plant meristems in which, in addition to the regular mitoses with the diploid chromosome number, occur also euploid or aneuploid chromosome number of certain chromosome within one individual (reviewed in D’amato, 1985). We tested the level of somatic aneuploidy in Lib male accession (termed also as aneusomaty – euploid or aneuploid chromosome number of certain chromosome within one individual). Basic chromosome number differed among the leaf tissue in one plant (2n; 2n + 2). We differentiated accessory chromosome 2 in metaphase (2n + 2) based on simultaneous localization of newly identified satellites Saaz293 and HuluTR120 (Fig. 4a) and 5S rDNA (Fig. S21). We observed variable levels of aneusomaty for chromosome 2 (two to three foci of Saaz293 and HuluTR120) in male interphase (Figs 4a, S22a). Interestingly, we detected aneusomaty only in male hop plants, while female individuals exhibited only even number of Saaz293, with minimal or no variability per plant (Fig. S22b).

**Fig. 4.**
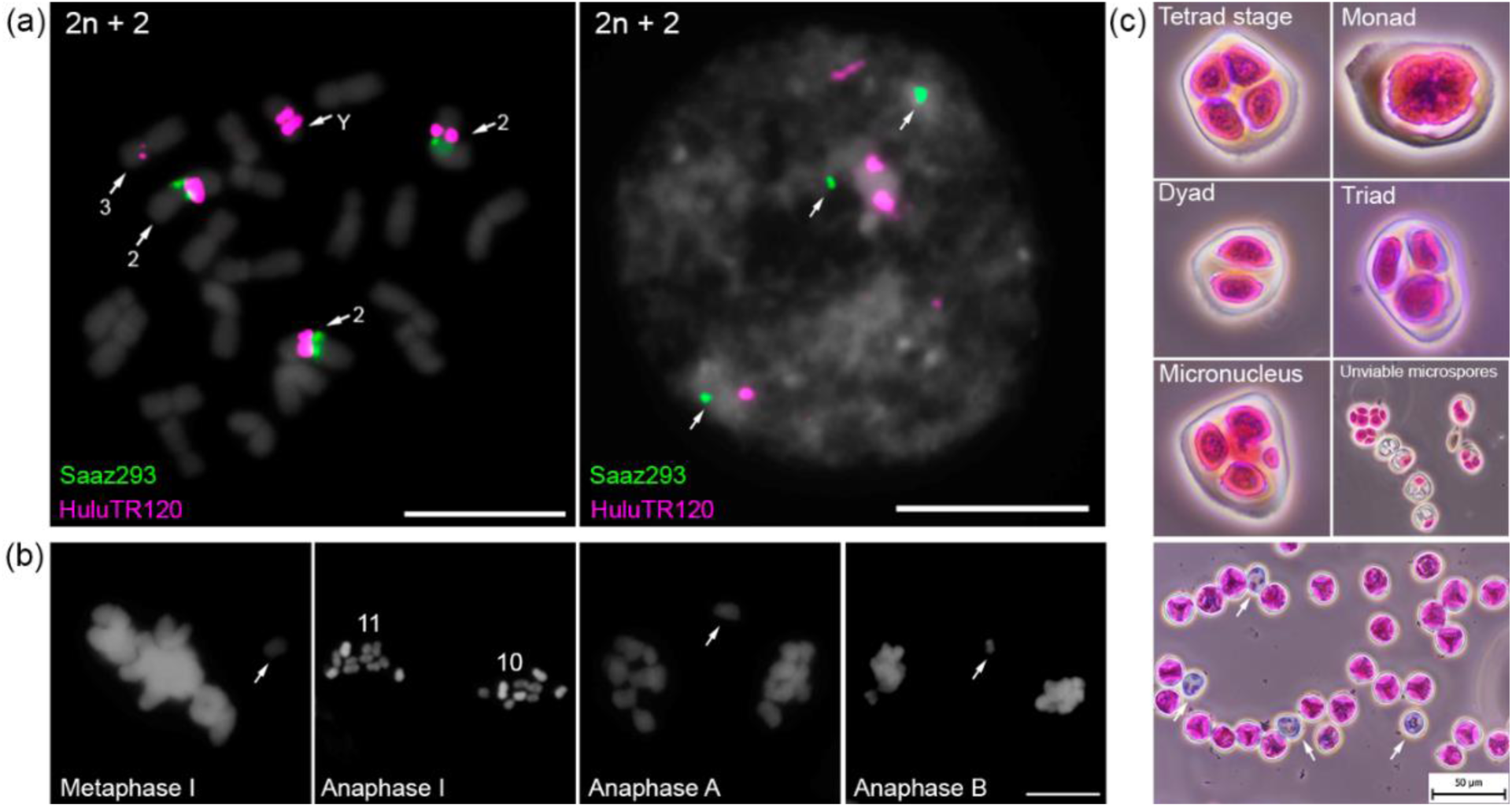
Effects of accessory chromosome 2 on mitosis, meiosis, and pollen grain development in male *Humulus lupulus*. (a) Aneuploidy (2n + 2) in male *H*. *lupulus* is characterized by three copies of chromosome 2 identified by signals of centromeric satellite Saaz293 (green) and HuluTR120 (magenta). Arrows indicate the three signals of satellite Saaz293 (green) providing direct evidence of aneusomaty in the male nucleus. Scale bar = 10 µm. (b) Observations of meiotic abnormalities include: an excluded bivalent from the metaphase plate (arrow), unbalanced segregation of homologous chromosomes to opposite cell poles, and lagging chromosomes during the anaphase A and anaphase B. Chromosomes and nuclei were counterstained with DAPI. Scale bar = 10 µm. (c) Comparison of regular tetrad stage with various form of unreduced pollen formation, including monad, dyad and triad. Presence of micronuclei and unviable microspores observed in three male *H*. *lupulus* plants. Viable pollen grains are stained purple, while nonviable pollen grains appear gray (arrows). The frequency of meiotic errors and unviable microspores is detailed in Tables S7, S8. Scale bar = 50 µm.

Based on previous observations of non-Mendelian inheritance in hop plants (Easterling *et al*., 2018, 2020), we screened PMCs at meiosis I (metaphase and anaphase) and meiosis II (tetrad stage). We found bivalents excluded from the metaphase plate during metaphase I, unbalanced segregation of homologous chromosomes to opposite poles of the cell, and the presence of lagging chromosomes in late anaphase I and telophase I (Fig. 4b). To assess the impact on microsporogenesis, we analyzed three *H*. *lupulus* males (Lib male, F_1_ progeny (Osvald’s clone 72 x Lib male), and wild hop). Alongside the balanced tetrad formation, we identified unreduced cells (monads, dyads, and triads) ranging from 5.36% to 9.24%, micronuclei (0.40-0.87%), and unviable microspores (2.50-6.96%) across the three different *Humulus* accessions, with a total of 10.57 to 12.72% meiotic errors (Fig. 4c, Table S7). Despite the number of meiotic abnormalities, pollen viability was significantly reduced only in Lib male accession (Fig. 4c, Table S7), harboring 3.95% of unviable pollen grains, compared to F1 male progeny (Osvald’s clone 72 x Lib male) and wild hop which had averages of 0.55% and 0.09% of unviable pollen grains, respectively (Table S8). These karyotypic observations and cytogenetic analysis indicate that the accessory chromosome 2 provides a direct link between the aneusomaty, meiotic errors, and decrease in fertility (Fig. 4).

## Discussion

Aberrant meiosis in plants can have both positive and negative consequences. Aside from aneuploidy, reduced fertility or sterility, altered seed development, and genetic instability, it also contributes to increases genetic diversity, promoting chromosomal rearrangement, and driving polyploidy. These processes make aberrant meiosis important force in promoting speciation and adaptation (reviewed in Zamariola *et al*., 2014; De Storme & Mason, 2014). Previous studies of *Humulus* genome revealed an unusual pattern of chromosomal segregation and non-Mendelian inheritance pattern during male meiosis in the North American hop variety (*H*. *lupulus* var. *neomexicanus*). The same study proposed that aberrant meiosis may be a natural feature, consequence of breeding of genetically divergent *Humulus* varieties or unusual centromere structure (Easterling *et al*., 2018, 2020). In this study, we assess whether the aberrant chromosome segregation is linked to unusual centromere organization, the existence of dicentric chromosome or other genomic factors.

### Structural features of centromeric landscape in *H*. *lupulus*

The centromeres of *H*. *lupulus* lack the higher-order repeats commonly found in most plants, such as *Cen178* in *Arabidopsis thaliana* (Wlodzimierz *et al*., 2023). ChIP-seq analysis using HlCENH3 demonstrates that *H*. *lupulus* centromeres of all chromosomes are primarily composed of SaazCRM1 CRM retrotransposons which are the major centrophilic TEs, and SaazCEN repeat (Figs 1b,c, 2b). The basic centromeric monomer unit in *H*. *lupulus* is 284 bp, with short subarrays of 39 bp subunits at the 3’SaazCEN terminus. Although the role of 39 bp subunits remains unclear, the average monomer length is consistent with those of other species which range from 23 bp in *Chionographis japonica* (Kuo *et al*., 2023) to 882 bp in *Pisum sativum* (Macas *et al*., 2023), and higher-order satellite array in *Vigna unguiculata* (Yang *et al*., 2023). The main characteristic feature of the centromeric repeat SaazCEN is its insertion into the centrophilic SaazCRM1, particularly within LTR domains (Fig. S19). Phylogenetic analysis of CRM retrotransposons revealed two categories: autonomous CRM elements, which have all canonical proteins and full ORF, and noaCRM elements. NoaCRMs exhibit heterogeneous structure and lack essential proteins for retrotransposition (Figs S16-S18). Given the sequence similarity in the PBS regions of both CRM categories, we propose that noaCRM relies on autonomous CRM for its function, as it also exhibits similarity to other shared canonical proteins. Our analysis estimates the insertion of noaCRM to be between 0.1 and 1.0 million years ago (Fig. S18). The insertion times of *Humulus* CRMs align with those of CRR elements in rice (Nagaki *et al*., 2005) and CRW elements in wheat (Liu *et al*., 2008). In wheat, the *Quinta* element represents a more recent insertion in both diploid and hexaploidy wheats compared to other CRW (Li *et al*., 2013). More recently, the insertion times of *RLG_Cereba* and *RLG_Quinta* chromovirus families in einkorn wheat were estimated to 0.0-1.0 Mya (Ahmed *et al*., 2023), closely matching the estimated insertion of centrophilic CRMs in *H*. *lupulus*, suggesting evolutionary parallels. These findings highlight the potential role of chromoviruses in maintaining centromere function and preserving its integrity, as reviewed in (Lisch, 2013; Naish & Henderson, 2024). Based on the sequencing data supported by ChIP-seq analysis, we identified two types of centromeres. The first type consists of SaazCEN and SaazCRM1 only (chromosomes 1, 4, 5, 7, and X), while the second centromere type additionally includes Saaz293, Saaz40, and Saaz85 (chromosomes 2, 3, 6 and 8). Physical localization of the latter three centromeric repeats confirmed their colocalization only on chromosomes 2 and 6. Notably, chromosome Y possesses HuluTR120, making this satellite a strong candidate for Y chromosome centromere function (Figs 1e,f, S7, S10). The presence of additional centromeric satellites is not surprising as most plant species typically have centromeres composed of several satellite tandem arrays (reviewed in Naish & Henderson, 2024). More importantly, Saaz40 and Saaz85 share approximately 50-60% sequence similarity, occupy the same positions and exhibit similar enrichment density on metaphase chromosomes (Figs S8, S13). This raises the question whether Saaz293, Saaz40, Saaz85, and HuluTR120 are potential new evolving centromeric sequences leading over time to the stabilization and refinement of centromeric repeats (reviewed in Naish & Henderson, 2024). As a rapid evolution of centromeric DNA sequences leads to high sequence divergence between closely related species (Lee *et al*., 2005), it makes the distribution of these centromeric repeats particularly unique, namely on chromosomes 2, 6, and Y (Figs 1a,d-g, 2b). However, it remains to be elucidated whether these repeats, and major CRMs together with SaazCEN, are shared with other species in the *Cannabaceae* family and other *Humulus* varieties. It is intriguing to speculate that the burst of CRM elements in *H*. *lupulus* centromeres may have occurred in the context of chromosomal rearrangements and genome evolution, particularly in the context of sex chromosome differentiation. In *Cannabis sativa*, for instance, the centromere of chromosome 7 is enriched with Harbinger TEs, with divergence time estimated at ∼20 Mya. Lynch *et al*. (2024) propose that these TEs contribute to genome rearrangements and centromeric evolution between *C. sativa* and *H. lupulus* (Lynch *et al*., 2024). Similarly, Zhang *et al*. (2023) identified centromeric satellite Hssat1 in *Humulus japonicus*, which is present in the centromeres of all chromosomes except for two Y chromosomes. The authors suggested that sex chromosomes in *H. japonicus* originated from centric fission events, potentially leading to the loss of centromeric-specific satellites on the Y chromosomes. In this study, however we did not find any similar sequences homologous to Hssat1. Although centromere evolution in related species remains largely unresolved, it is becoming apparent that the two Y chromosomes in *H. japonicus*, which lack major centromeric repeats, may illustrate a case of genomic convergence in centromere evolution comparable to *H*. *lupulus*. Given the relative recent insertion of CRMs elements in the centromeric regions in this work, we anticipate that these elements in *H*. *lupulus* and potentially in *H*. *japonicus* Y chromosomes, may play a key role in maintaining centromere integrity, similarly as in einkorn wheat (Ahmed *et al*., 2023). Therefore, a comprehensive understanding of centromere organization in closely related species, such as *C. sativa* and *H. japonicus*, will shed light on the evolutionary dynamics of centromere structure and function. As mentioned above, the chromosome 2, is enriched for Saaz293 satellite array and pericentromeric localization of HuluTR120 (Fig. 1d). This chromosome exhibits a higher-order array structure compared to other autosomes, including chromosomes 6 and 8 (Figs 1, S4). Remarkably, chromosome 2 shows low enrichment for both major centromeric repeats, SaazCEN and SaazCRM1 (Fig. S5). A similar centromere composition has been reported in sunflower, where all chromosomes are enriched with LINE elements, with the exception of one chromosome that harbors centromeric satellite (Nagaki *et al*., 2005). This leads us, to speculate that the Saaz293 may diverged from the last common ancestor, hinting the potential for neocentromere formation. A similar scenario has been proposed for the unique centromeric landscape in potato which has two types of centromeres (Gong *et al*., 2012). While the origin of newly identified repeats remains unclear, our findings indicate that chromosome 2 is likely involved in the non-Mendelian segregation patterns previously reported by Easterling *et al*. (2018, 2020).

### Non-Mendelian segregation patterns and the uniqueness of chromosome 2

We detected aneuploidy (2n + 2) with one accessory chromosome 2 in Lib male accession, consistent with previously reported meiotic abnormalities, and identified these metaphases based on Saaz293 and HuluTR120 distribution (Fig. 4a). Our results provide a direct link between the unique centromere organization of chromosome 2 and its centromere role in chromosomal segregation during cell division. At the same time, by screening of PMCs during meiosis I and II in Lib male, F_1_ progeny (Osvald’s clone 72 x Lib male), and wild hop, we confirmed its aberrant chromosomal segregation (Fig. 4b). This aberrant segregation results in unviable microspores, varying severity of unreduced tetrad formation, and the presence of micronuclei, all of which contribute to reduced pollen viability. These observations corroborate the non-Mendelian ratio of 5S rDNA at the tetrad stage (Easterling *et al*., 2018), as both satellites Saaz293 and HuluTR120 are localized on the same chromosome, which also harbors the 5S rDNA subunit on chromosome 2. Despite the limited FISH DNA markers prevented identification of the second accessory chromosome in metaphase (2n + 2), we hypothesize that the second chromosome may originate from homologous chromosome pair 3 which harbors HuluTR120 only within one chromosome of this pair (Figs 1e,f, S11d,e). To test this hypothesis, unique DNA oligo painting probes will need to be developed as previously described in *Silene latifolia* (Bačovský *et al*., 2020) or reviewed in Jiang (2019) and Hobza *et al*. (2024).

Interestingly, the utility of Saaz293 lies in its unique chromosomal localization (compared to 5S rDNA or HuluTR120) that allows easy tracking of non-Mendelian segregation patterns across various hop accessions in future studies. In combination with techniques like high-content imaging and cell population screening (Hobza *et al*., 2024), Saaz293 could serve as a promising marker for hop breeding, allowing to test the tissue cultures stability and monitor spontaneous chromosome instability during *in vitro* cultivation (Abugammie *et al*., 2024). In the context of previous studies, we hypothesize that non-Mendelian segregation patterns result from either nondisjunction (Fig. 5a) of two chromosome pairs or incorrect microtubule attachment during metaphase (Fig. 5b). Both scenarios may result in the formation of aneuploid cells, monads, dyads, triads, and micronuclei, collectively reducing pollen viability (Fig. 5c). We propose that the unique centromere structure of chromosome 2 plays an crucial role in chromosome segregation and the observed non-Mendelian segregation patterns (Fig. 4b,c).

**Fig. 5.**
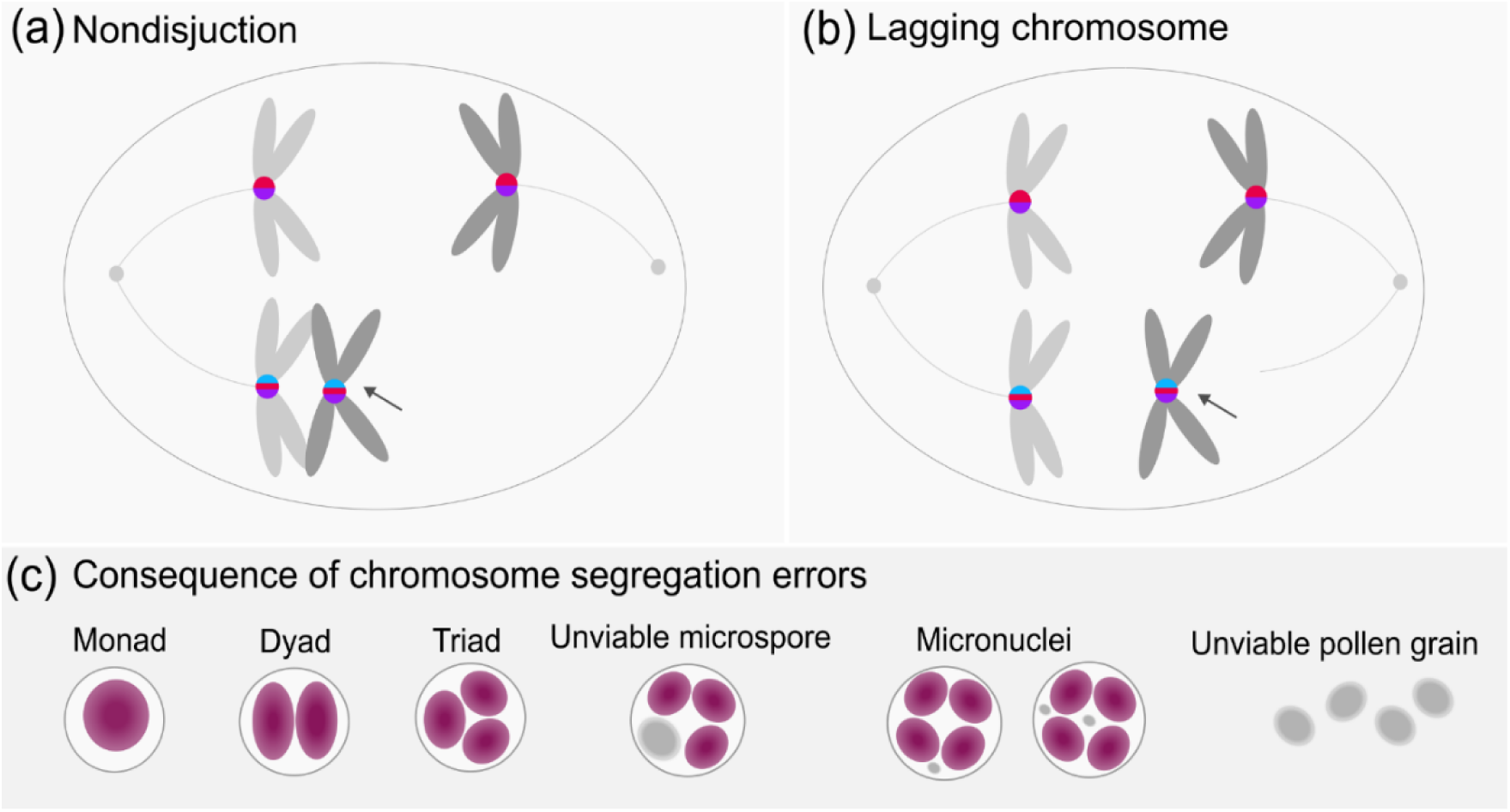
Schematic model of chromosome 2 aberrant segregation in *Humulus lupulus*. (a) Nondisjunction and (b) lagging chromosome 2 (arrows) result in the non-Mendelian segregation patterns (c) associated with microspore reduction, micronuclei formation and lower pollen viability.

This study provides the first detailed survey on the centromeric landscape of dioecious plant *H*. *lupulus* and provide direct evidence between unique centromeric structure of chromosome 2 and previously published non-Mendelian segregation patterns. The precise localization of centromeric repeats SaazCEN and SaazCRM1 on Y chromosome in the Lib male further refines the position of HSR1 subtelomeric probe (Fig. S10) than was previously described by Divashuk *et al*., (2011). The HSR1 positive regions are located in the subtelomeric region on the p-arm of Y chromosome and in both the pericentromeric and subtelomeric regions on the X chromosome. These results posited the pseudoautosomal region (PAR) on the p-arm of the Y chromosome. This is further supported by the self-GISH approach highlighting the male-specific region and excluding the PAR at the termini of Y chromosome p-arm (Razumova *et al*., 2023). Additionally, our results enable the testing of other male accessions for non-Mendelian segregation and pave the way for understanding the evolutionary processes that led to species divergence within the *Cannabaceae* family.

## Acknowledgements

We would like to thank Alena Henychova from Hop Research Institute in Žatec (Czech Republic) for her valuable contribution to hop cultivation. Hop breeding was conducted as part of the collection of hop genetic resources, under the project MZe - 51834/2017-MZE-17253/6.2.1, titled “National program for the conservation and the use of genetic resources of plants and agrobiodiversity”, founded by the Ministry of Agriculture. Computational resources were provided by the e-INFRA CZ project (ID:90254), supported by the Ministry of Education, Youth and Sports of the Czech Republic and by the ELIXIR-CZ project (ID:90255), part of the international ELIXIR infrastructure. This work was supported by the Czech Science Foundation grant no.: 22-00301S and from the project TowArds Next GENeration Crops, reg. no. CZ.02.01.01/00/22_008/0004581 of the ERDF Programme Johannes Amos Comenius.

## Competing interests

The authors declare no competing interests.

## Author contributions

VB, LH, VH, JS, and RH planned and designed research. LH, VB, and PN performed experiments. RČ and PJ analyzed the data. LH and VB wrote the main text of the manuscript. HT, AT, and TI performed genome sequence and assembly of *Humulus* plants. TA and EO provided sequencing data. JP provided plant material. All authors read and approved the final version of the manuscript.

## Data availability

All sequencing data are available in the European Nucleotide Archive (ENA) under accession number PRJEB81858. Saaz hop female reference genome is accessible at AP031542-AP031551, AP035121-AP035448.

---

**Fig. S1** CENH3 blast-n in *Humulus lupulus* and multiple alignment of HlCENH3 sequence with other plant species.

**Fig. S2** The sequence variability of HlCENH3 gene in *Humulus lupulus*.

**Fig. S3** The localization of HlCENH3 antibody in interphase nuclei of Saaz female *Humulus lupulus*.

**Fig. S4** Centromere repeat array characterization for each chromosome in Saaz female *H*. *lupulus* genome.

**Fig. S5** Major centromeric repeat arrays for each chromosome in Saaz female *H*. *lupulus* genome.

**Fig. S6** Male karyotype of *Humulus lupulus*, 2n = 20, XY.

**Fig. S7** Detailed analysis of chromosomes 3 and Y in three different *Humulus lupulus* male accessions.

**Fig. S8** The localization of centromeric satellite Saaz40 and 45S rDNA on chromosome 6 in male and female of *Humulus lupulus*.

**Fig. S9** The localization of centromeric satellite Saaz85 and 45S rDNA on chromosome 6 in male and female *Humulus lupulus*.

**Fig. S10** Chromosome pairing and organization during diakinesis of *Humulus lupulus* Lib male.

**Fig. S11** Distribution of major centromeric repeats on metaphase chromosomes of *Humulus lupulus* Saaz female.

**Fig. S12** Distribution of major centromeric repeats on metaphase chromosomes of *Humulus lupulus* Lib male.

**Fig. S13** Dot plot and sequence similarity of three major centromeric satellites Saaz85, Saaz293 and Saaz40 of chromosomes 2, 3, 6, and 8.

**Fig. S14** LTR retrotransposons composition of all centromeres of *Humulus lupulus* Saaz female.

**Fig. S15** Distribution and estimated insertion time of Ty1/*Copia* and Ty3/*Gypsy* families of LTR retrotransposons for all chromosomes.

**Fig. S16** The insertion time of autonomous, dominant non-autonomous, and minor non-autonomous CRMs.

**Fig. S17** The full-length distribution of centromeric Ty3/*Gypsy* CRM retrotransposons in *Humulus lupulus*.

**Fig. S18** Phylogenetic analysis of the autonomous and non-autonomous CRM retrotransposons correlated with insertion age (MYA).

**Fig. S19** Distribution of SaazCEN and SaazCRM1 within the autonomous and non-autonomous CRM retrotransposons in *Humulus* centromeres.

**Fig. S20** Localization of HlCENH3 within CRM retrotransposons in *Humulus* centromeres.

**Fig. S21** Localization of the major centromeric satellites on chromosomes 2 and 6.

**Fig. S22** Positioning of chromosome 2 within an interphase nucleus of male and female *Humulus lupulus*.

**Table S1** List of *Humulus lupulus* plants utilized in this study.

**Table S2** List of primers used for PCR and preparation of FISH probes.

**Table S3** Enzyme mixture used for digestion of young leaves.

**Table S4** Clusters of tandem repeats selected for FISH including ChipSeq Mapper outputs.

**Table S5** Genomic fraction of repetitive DNA in the *Humulus lupulus* Saaz and Lib male genome.

**Table S6** Tandem repeats in *H*. *lupulus* Saaz and Lib male genome identified by RepeatEplorer2 pipeline.

**Table S7** Number and frequency of meiotic abnormalities in three male accessions of *Humulus lupulus*.

**Table S8** Number and frequency of viable and unviable pollen grain in three male accessions of *Humulus lupulus*.

**Methods S1** Extraction of DNA, Genome sequencing and Characterization of repetitive DNA.

**Notes S1** Repeatome analysis of Saaz female and Lib male of *H*. *lupulus*.

## Notes

### Competing Interest Statement

The authors have declared no competing interest.

## References

Abugammie B, Wang R, Hu Y, Pang J, Luan Y, Liu B, Jiang L, Lv R. 2024. Spontaneous chromosome instability and tissue culture-induced karyotypic alteration in wheat–Thinopyrum intermedium alien addition lines. Planta 260: 17.

Ahmed HI, Heuberger M, Schoen A, Koo DH, Quiroz-Chavez J, Adhikari L, Raupp J, Cauet S, Rodde N, Cravero C, et al. 2023. Einkorn genomics sheds light on history of the oldest domesticated wheat. Nature 620: 830–838.

Bačovský V, Čegan R, Šimoníková D, Hřibová E, Hobza R. 2020. The Formation of Sex Chromosomes in Silene latifolia and S. dioica Was Accompanied by Multiple Chromosomal Rearrangements. Frontiers in Plant Science 11.

Bačovský V, Houben A, Kumke K, Hobza R. 2019. The distribution of epigenetic histone marks differs between the X and Y chromosomes in Silene latifolia. Planta 250: 487–494.

Bi G, Zhao S, Yao J, Wang H, Zhao M, Sun Y, Hou X, Haas FB, Varshney D, Prigge M, et al. 2024. Near telomere-to-telomere genome of the model plant Physcomitrium patens. Nature Plants 10: 327–343.

Bolger AM, Lohse M, Usadel B. 2014. Trimmomatic: A flexible trimmer for Illumina sequence data. Bioinformatics 30: 2114–2120.

D’amato F. 1985. Cytogenetics of plant cell and tissue cultures and their regenerates. Critical Reviews in Plant Sciences 3: 73–112.

De Storme N, Mason A. 2014. Plant speciation through chromosome instability and ploidy change: Cellular mechanisms, molecular factors and evolutionary relevance. Current Plant Biology 1: 10–33.

Divashuk MG, Alexandrov OS, Kroupin PY, Karlov GI. 2011. Molecular cytogenetic mapping of Humulus lupulus sex chromosomes. Cytogenetic and Genome Research 134: 213–219.

Draizen EJ, Shaytan AK, Mari∼ No-Ram Irez L, Talbert PB, Landsman D, Panchenko AR. 2016. HistoneDB 2.0: a histone database with variants-an integrated resource to explore histones and their variants. Database 2016: 14–14.

Easterling KA, Pitra NJ, Jones RJ, Lopes LG, Aquino JR, Zhang D, Matthews PD, Bass HW. 2018. 3D molecular cytology of hop (humulus lupulus) meiotic chromosomes reveals non-disomic pairing and segregation, aneuploidy, and genomic structural variation. Frontiers in Plant Science 871.

Easterling KA, Pitra NJ, Morcol TB, Aquino JR, Lopes LG, Bussey KC, Matthews PD, Bass HW. 2020. Identification of tandem repeat families from long-read sequences of Humulus lupulus. PLoS ONE 15.

Gent JI, Wang N, Dawe RK. 2017. Stable centromere positioning in diverse sequence contexts of complex and satellite centromeres of maize and wild relatives. Genome Biology 18.

Gong Z, Wu Y, Koblížková A, Torres GA, Wang K, Iovene M, Neumann P, Zhang W, Novák P, Robin Buell C, et al. 2012. Repeatless and repeat-based centromeres in potato: Implications for centromere evolution. Plant Cell 24: 3559–3574.

Haunold A. 1974. Meiotic Chromosome Behavior and Pollen Fertility of a Triploid Hop 1. Crop Science 14: 849–852.

Henikoff S, Ahmad K, Malik HS. 2001. The Centromere Paradox: Stable Inheritance with Rapidly Evolving DNA. Science 293: 1098–1102.

Heuberger M, Koo D-H, Ahmed HI, Tiwari VK, Abrouk M, Poland J, Krattinger SG, Wicker T. 2024. Evolution of Einkorn wheat centromeres is driven by the mutualistic interplay of two LTR retrotransposons. Mobile DNA 15: 16.

Hill ST, Sudarsanam R, Henning J, Hendrix D. 2017. HopBase: A unified resource for Humulus genomics. Database 2017.

Hobza R, Bačovský V, Čegan R, Horáková L, Hubinský M, Janíček T, Janoušek B, Jedlička P, Kružlicová J, Kubát Z, et al. 2024. Sexy ways: approaches to studying plant sex chromosomes. Journal of Experimental Botany.

Houben A, Demidov D, Gernand D, Meister A, Leach CR, Schubert I. 2003. Methylation of histone H3 in euchromatin of plant chromosomes depends on basic nuclear DNA content. Plant Journal 33: 967–973.

Houben A, Schroeder-Reiter E, Nagaki K, Nasuda S, Wanner G, Murata M, Endo TR. 2007. CENH3 interacts with the centromeric retrotransposon cereba and GC-rich satellites and locates to centromeric substructures in barley. Chromosoma 116: 275–283.

Jiang J. 2019. Fluorescence in situ hybridization in plants: recent developments and future applications. Chromosome Research 27: 153–165.

Katoh K, Standley DM. 2013. MAFFT multiple sequence alignment software version 7: Improvements in performance and usability. Molecular Biology and Evolution 30: 772–780.

Kuo YT, Câmara AS, Schubert V, Neumann P, Macas J, Melzer M, Chen J, Fuchs J, Abel S, Klocke E, et al. 2023. Holocentromeres can consist of merely a few megabase-sized satellite arrays. Nature Communications 14.

Kyte J, Doolittle RF. 1982. A simple method for displaying the hydropathic character of a protein. Journal of Molecular Biology 157: 105–132.

Langdon T, Seago C, Mende M, Leggett M, Thomas H, Forster JW, Jones RN, Jenkins G. 2000. Retrotransposon evolution in diverse plant genomes. Genetics 156: 313–325.

Lee H-R, Zhang W, Langdon T, Jin W, Yan H, Cheng Z, Jiang J. 2005. Chromatin immunoprecipitation cloning reveals rapid evolutionary patterns of centromeric DNA in Oryza species. Proceedings of the National Academy of Sciences 102: 11793–11798.

Li B, Choulet F, Heng Y, Hao W, Paux E, Liu Z, Yue W, Jin W, Feuillet C, Zhang X. 2013. Wheat centromeric retrotransposons: The new ones take a major role in centromeric structure. Plant Journal 73: 952–965.

Lisch D. 2013. How important are transposons for plant evolution? Nature Reviews Genetics 14: 49–61.

Liu Y, Su H, Pang J, Gao Z, Wang X-J, Birchler JA, Han F. 2015. Sequential de novo centromere formation and inactivation on a chromosomal fragment in maize. Proceedings of the National Academy of Sciences 112.

Liu Z, Yue W, Li D, Wang RRC, Kong X, Lu K, Wang G, Dong Y, Jin W, Zhang X. 2008. Structure and dynamics of retrotransposons at wheat centromeres and pericentromeres. Chromosoma 117: 445–456.

Lunerová J, Vozárová R. 2023. Preparation of Male Meiotic Chromosomes for Fluorescence In Situ Hybridization and Immunodetection with Major Focus on Dogroses. In: Heitkam T, Garcia S, eds. Methods in Molecular Biology. Plant Cytogenetics and Cytogenomics. New York, NY: Springer US, 337–349.

Lynch RC, Padgitt-Cobb LK, Garfinkel AR, Knaus BJ, Hartwick NT, Allsing N, Aylward A, Mamerto A, Kitony JK, Colt K, et al. 2024. Domesticated cannabinoid synthases amid a wild mosaic cannabis pangenome. bioRxiv: 2024.05.21.595196-2024.05.21.595196.

Macas J, Robledillo LÁ, Kreplak J, Novák P, Koblížková A, Vrbová I, Burstin J, Neumann P. 2023. Assembly of the 81.6 Mb centromere of pea chromosome 6 elucidates the structure and evolution of metapolycentric chromosomes. PLoS Genetics 19.

Maheshwari S, Ishii T, Brown CT, Houben A, Comai L. 2017. Centromere location in Arabidopsis is unaltered by extreme divergence in CENH3 protein sequence. Genome Research 27: 471–478.

Majka J, Glombik M, Doležalová A, Kneřová J, Ferreira MTM, Zwierzykowski Z, Duchoslav M, Studer B, Doležel J, Bartoš J, et al. 2023. Both male and female meiosis contribute to non-Mendelian inheritance of parental chromosomes in interspecific plant hybrids (Lolium × Festuca). New Phytologist 238: 624–636.

Mandáková T, Hloušková P, Koch MA, Lysak MA. 2020. Genome Evolution in Arabideae Was Marked by Frequent Centromere Repositioning. The Plant Cell 32: 650–665.

McAdam EL, Freeman JS, Whittock SP, Buck EJ, Jakse J, Cerenak A, Javornik B, Kilian A, Wang CH, Andersen D, et al. 2013. Quantitative trait loci in hop (Humulus lupulus L.) reveal complex genetic architecture underlying variation in sex, yield and cone chemistry. BMC Genomics 14.

Melters DP, Bradnam KR, Young HA, Telis N, May MR, Ruby JG, Sebra R, Peluso P, Eid J, Rank D, et al. 2013. Comparative analysis of tandem repeats from hundreds of species reveals unique insights into centromere evolution. Genome Biology 14.

Mendiburo MJ, Padeken J, Fülöp S, Schepers A, Heun P. 2011. Drosophila CENH3 Is Sufficient for Centromere Formation. Science 334: 683–686.

Nagaki K, Neumann P, Zhang D, Ouyang S, Buell CR, Cheng Z, Jiang J. 2005. Structure, divergence, and distribution of the CRR centromeric retrotransposon family in rice. Molecular Biology and Evolution 22: 845–855.

Naish M, Henderson IR. 2024. The structure, function, and evolution of plant centromeres. Genome Research 34.

Natsume S, Takagi H, Shiraishi A, Murata J, Toyonaga H, Patzak J, Takagi M, Yaegashi H, Uemura A, Mitsuoka C, et al. 2015. The Draft Genome of Hop (Humulus lupulus), an Essence for Brewing. Plant and Cell Physiology 56: 428–441.

Navrátilová P, Toegelová H, Tulpová Z, Kuo YT, Stein N, Doležel J, Houben A, Šimková H, Mascher M. 2022. Prospects of telomere-to-telomere assembly in barley: Analysis of sequence gaps in the MorexV3 reference genome. Plant Biotechnology Journal 20: 1373–1386.

Neumann P, Navrátilová A, Schroeder-Reiter E, Koblížková A, Steinbauerová V, Chocholová E, Novák P, Wanner G, Macas J. 2012. Stretching the rules: Monocentric chromosomes with multiple centromere domains. PLoS Genetics 8.

Neumann P, Novák P, Hoštáková N, Macas J. 2019. Systematic survey of plant LTR-retrotransposons elucidates phylogenetic relationships of their polyprotein domains and provides a reference for element classification. Mobile DNA 10.

Neumann P, Schubert V, Fuková I, Manning JE, Houben A, Macas J. 2016. Epigenetic Histone Marks of Extended Meta-Polycentric Centromeres of Lathyrus and Pisum Chromosomes. Frontiers in Plant Science 7.

Neve RA. 1958. Sex Chromosomes in the Hop Humulus lupulus. Nature 181: 1084–1085.

Neve RA. 1991. Hops. Dordrecht: Springer Netherlands.

Novák P, Neumann P, Macas J. 2020. Global analysis of repetitive DNA from unassembled sequence reads using RepeatExplorer2. Nature Protocols 15: 3745–3776.

Novák P, Neumann P, Pech J, Steinhaisl J, MacAs J. 2013. RepeatExplorer: A Galaxy-based web server for genome-wide characterization of eukaryotic repetitive elements from next-generation sequence reads. Bioinformatics 29: 792–793.

Ono T. 1955. Studies in Hop. Chromosomes of Common Hop and its relatives. Bulletin of Brewing Science: 3–65.

Padgitt-Cobb LK, Pitra NJ, Matthews PD, Henning JA, Hendrix DA. 2023. An improved assembly of the “Cascade” hop (*Humulus lupulus*) genome uncovers signatures of molecular evolution and refines time of divergence estimates for the Cannabaceae family. Horticulture Research 10: uhac281.

Peterson R, Slovin JP, Chen C. 2010. A simplified method for differential staining of aborted and non-aborted pollen grains. International Journal of Plant Biology 1: 66–69.

Price MN, Dehal PS, Arkin AP. 2010. FastTree 2 - Approximately maximum-likelihood trees for large alignments. PLoS ONE 5.

Razumova OV, Divashuk MG, Alexandrov OS, Karlov GI. 2023. GISH painting of the Y chromosomes suggests advanced phases of sex chromosome evolution in three dioecious Cannabaceae species (Humulus lupulus, H. japonicus, and Cannabis sativa). Protoplasma 260: 249–256.

Sacchi B, Humphries Z, Kružlicová J, Bodláková M, Pyne C, Choudhury BI, Gong Y, Bačovský V, Hobza R, Barrett SCH, et al. 2024. Phased Assembly of Neo-Sex Chromosomes Reveals Extensive Y Degeneration and Rapid Genome Evolution in Rumex hastatulus (M Wilson, Ed.). Molecular Biology and Evolution 41: msae074.

Sanei M, Pickering R, Kumke K, Nasuda S, Houben A. 2011. Loss of centromeric histone H3 (CENH3) from centromeres precedes uniparental chromosome elimination in interspecific barley hybrids. Proceedings of the National Academy of Sciences of the United States of America 108.

Seefelder S, Ehrmaier H, Schweizer G, Seigner E. 2000. Male and female genetic linkage map of hops, Humulus lupulus. Plant Breeding 119: 249–255.

Sinotô Y. 1929. On the Tetrapartite Chromosome in Humulus Lupulus. Proceedings of the Imperial Academy 5: 46–47.

Small E. 1978. A Numerical and Nomenclatural Analysis of Morpho-Geographic Taxa of Humulus. Systematic Botany 3: 37.

Tarailo-Graovac M, Chen N. 2009. Using RepeatMasker to Identify Repetitive Elements in Genomic Sequences. Current Protocols in Bioinformatics 25.

Vasimuddin Md, Misra S, Li H, Aluru S. 2019. Efficient Architecture-Aware Acceleration of BWA-MEM for Multicore Systems. In: 2019 IEEE International Parallel and Distributed Processing Symposium (IPDPS). Rio de Janeiro, Brazil: IEEE, 314–324.

Winge O. 1923. On sex chromosomes, sex determination and preponderance of females in some dioecious plants. Comptes-rendus des travaux du Laboratoire de Carlsberg 15: 1–16.

Wlodzimierz P, Rabanal FA, Burns R, Naish M, Primetis E, Scott A, Mandáková T, Gorringe N, Tock AJ, Holland D, et al. 2023. Cycles of satellite and transposon evolution in Arabidopsis centromeres. Nature 618: 557–565.

Yang Y, Wu Z, Wu Z, Li T, Shen Z, Zhou X, Wu X, Li G, Zhang Y. 2023. A near-complete assembly of asparagus bean provides insights into anthocyanin accumulation in pods. Plant Biotechnology Journal 21: 2473–2489.

Zamariola L, Tiang CL, De Storme N, Pawlowski W, Geelen D. 2014. Chromosome segregation in plant meiosis. Frontiers in Plant Science 5: 279.

Zanoli P, Zavatti M. 2008. Pharmacognostic and pharmacological profile of Humulus lupulus L. Journal of Ethnopharmacology 116: 383–396.

Zhang D, Easterling KA, Pitra NJ, Coles MC, Buckler ES, Bass HW, Matthews PD. 2017. Non-Mendelian Single-Nucleotide Polymorphism Inheritance and Atypical Meiotic Configurations are Prevalent in Hop. The Plant Genome 10.

Zhang L, Hu J, Han X, Li J, Gao Y, Richards CM, Zhang C, Tian Y, Liu G, Gul H, et al. 2019. A high-quality apple genome assembly reveals the association of a retrotransposon and red fruit colour. Nature Communications 10: 1494.

Zhang GJ, Jia KL, Wang J, Gao WJ, Li SF. 2023. Genome-wide analysis of transposable elements and satellite DNA in Humulus scandens, a dioecious plant with XX/XY1Y2 chromosomes. Frontiers in Plant Science 14.

Zhang Y, Liu T, Meyer CA, Eeckhoute J, Johnson DS, Bernstein BE, Nussbaum C, Myers RM, Brown M, Li W, et al. 2008. Model-based analysis of ChIP-Seq (MACS). Genome Biology 9.

Zhao J, Xie Y, Kong C, Lu Z, Jia H, Ma Z, Zhang Y, Cui D, Ru Z, Wang Y, et al. 2023. Centromere repositioning and shifts in wheat evolution. Plant Communications 4: 100556.

