## Supplemental Files for "Centromeric repeat diversity underlies non-Mendelian segregation pattern in hop (*Humulus lupulus*)"

**Fig. S1** CENH3 blast-n in *Humulus lupulus* and multiple alignment of HlCENH3 sequence with other plant species. The position of the selected peptide for polyclonal antibody of HlCENH3 is shown in pink, the position of other histone domains - alphaN, alpha1, alpha2, alpha3 and linker are shown in orange, green and light green, respectively.

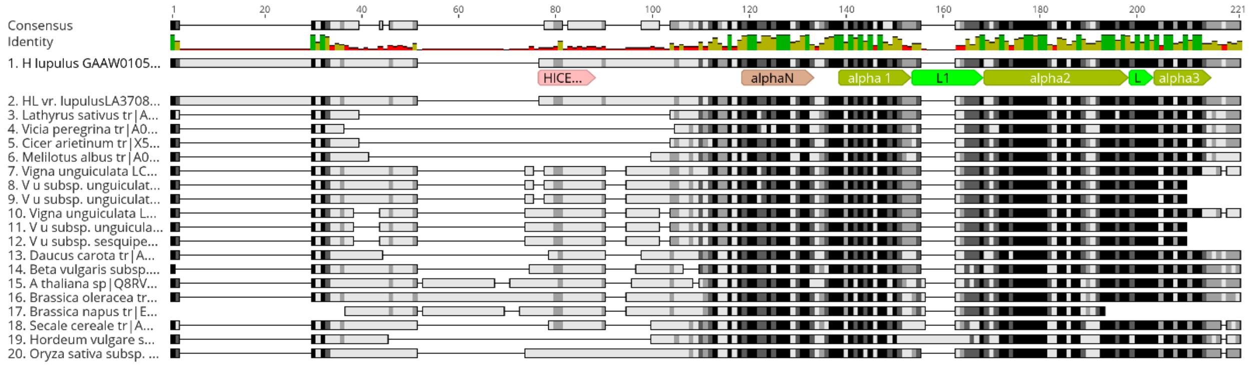

**Fig. S2** The sequence variability of HlCENH3 gene in *Humulus lupulus*. Eight sequences were amplified from colony PCR. The HlCENH3 gene display little or no variability within the *H. lupulus* genome.

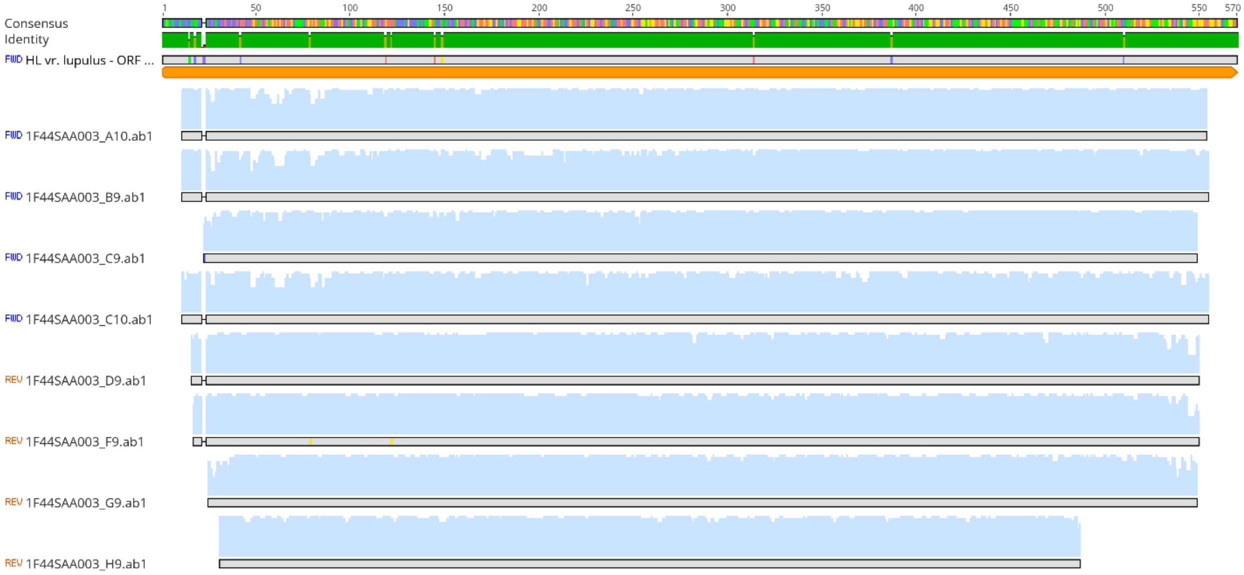

**Fig. S3** The localization of HlCENH3 antibody in interphase nuclei of Saaz female *Humulus lupulus*. Immunostaining with HlCENH3 displays distinct signals (green) in interphase nuclei. Nuclei were derived from two independent experiments. Interphase nuclei were counterstained with DAPI. Scale bar = 10 µm.

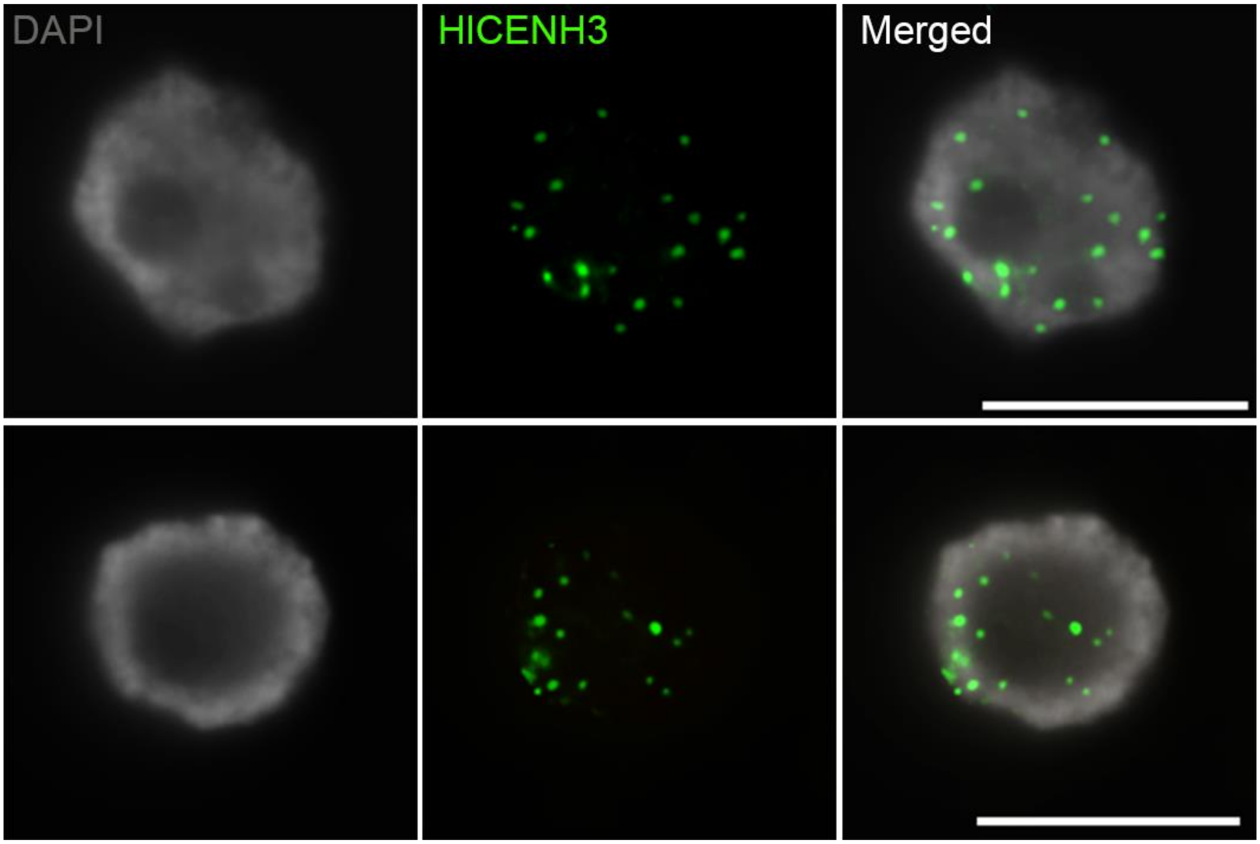

**Fig. S4** Centromere repeat array characterization for each chromosome in Saaz female *H*. *lupulus* genome. Dotplot for each centromere reveals high-order repeat arrays that are made by colored percent identity. (a) First three lines show HlCENH3 ChIP-seq data compared to input control. ChIP-seq data are shown in duplicate HlChip_A (green) and HlChip_B (blue). (b) The second three lines show the distribution of autonomous and non-autonomous CRMs, Retand and Tekay in centromeric regions. Heatmaps display pairwise sequence identity for centromere of each chromosome.

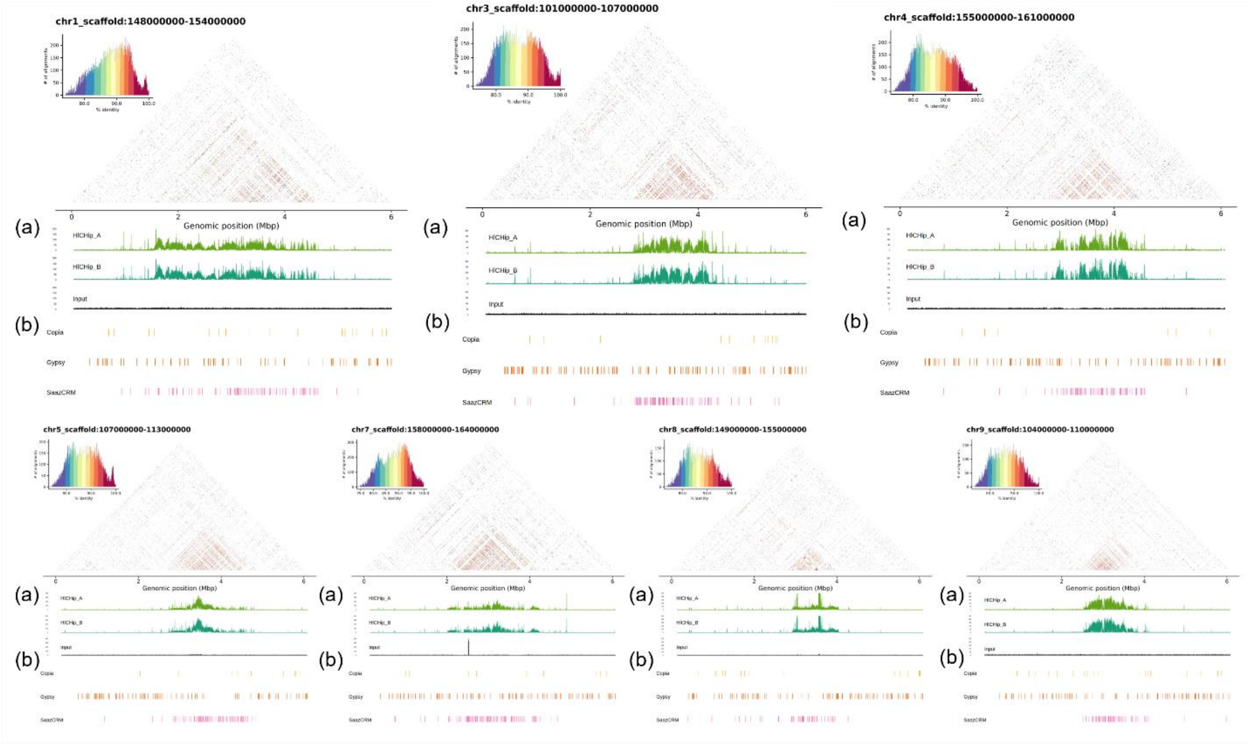

**Fig. S5** Major centromeric repeat arrays for each chromosome in Saaz female *H. lupulus* genome. First line shows HICENH3 binding region domain. The second line shows the distribution of major centromeric-specific repeats for each chromosome. Main centromeric repeat SaazCEN and centromere specific retrotransposon SaazCRM1, were observed to be shared among all chromosomes. Note specific distribution of three satellites Saaz40, Saaz85, and Saaz293 on chromosomes 2, 3, 6, and 8. Interestingly, chromosome 2 possess two HICENH3 positive regions which align with the distribution of chromosome 2-specific centromeric satellites.

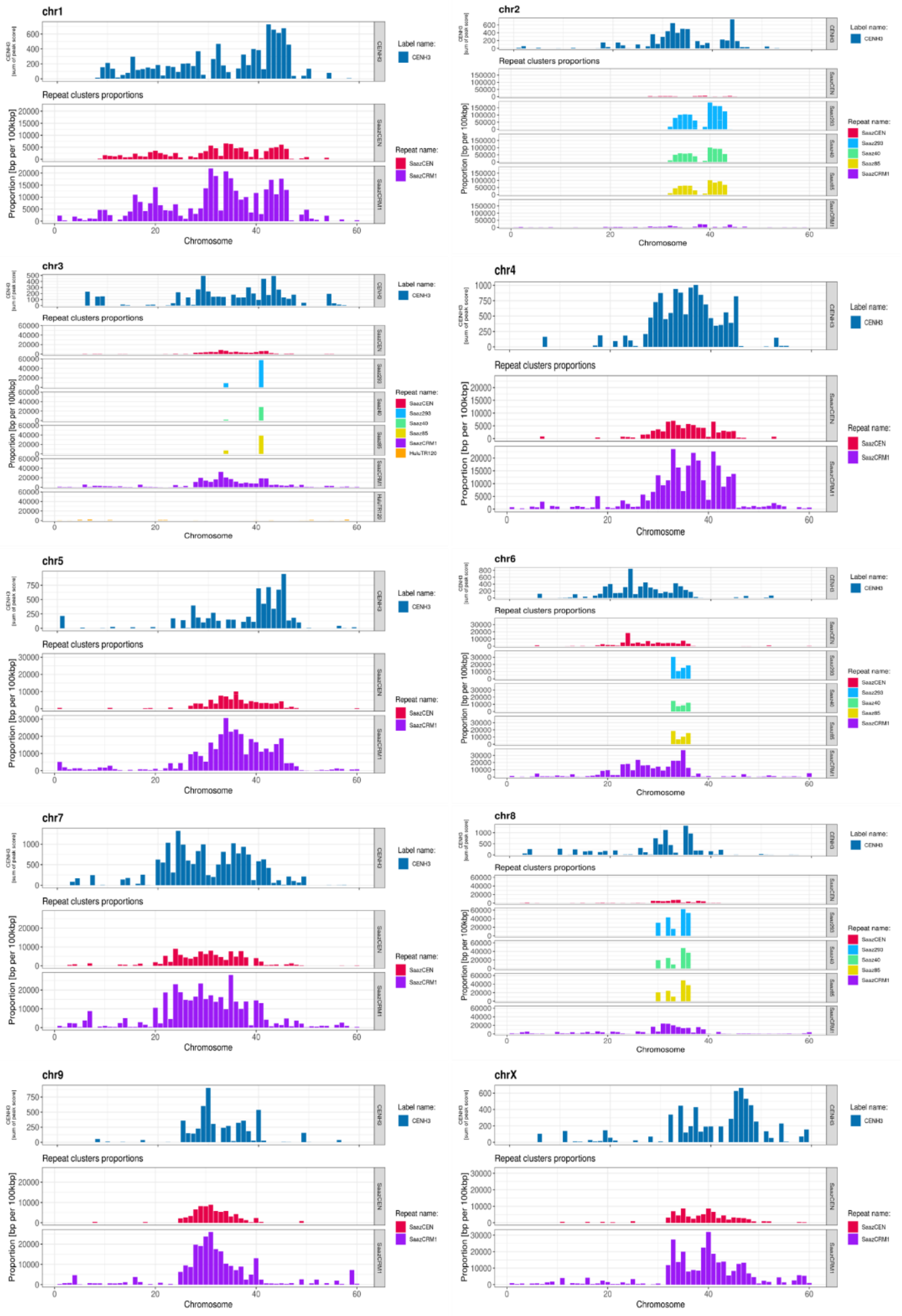

**Fig. S6** Male karyotype of *Humulus lupulus*, 2n = 20, XY. Centromeric repeat SaazCEN (red; newly identified in this study) and previously published subtelomeric repeat HSR (cyan) were used to distinguish p- and q-arms, and X and Y chromosomes. Satellite HSR is localized in the subtelomeric region of all autosomes, except chromosome 6, and p-arm of X chromosome (HSR signal in pericentromeric region). Y chromosome displays HSR satellite only on the p-arm in subtelomeric region. Chromosome 6 is missing the HSR satellite on p-arm. Mitotic chromosomes were counterstained with DAPI. Scale bar = 10 μm.

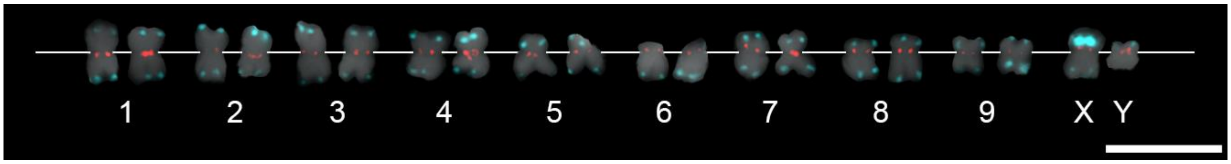

**Fig. S7** Detailed analysis of chromosomes 3 and Y in three different *Humulus lupulus* male accessions. Three satellites were used to differentiate chromosomes 2, 3 and Y – pericentromeric HuluTR120 (magenta), chromosome 2 centromere-specific Saaz293 (green) and subtelomeric HSR (cyan) in accessions (a) 15246, (b) 15249 and (c) 15276. Male accession 15246 (a) possesses locus of HuluTR120 only within one chromosome of autosomal pair 3. Compared to 15246, accessions 15249 (b) and 15276 (c) have even number of loci for HuluTR120. Therefore, accession 15246 represents hybrid constitution of chromosome 3 of unknown origin. Two distinct loci of HuluTR120 on chromosome Y are present in all studied accessions. Chromosomes were counterstained with DAPI. Scale bar = 10 μm.

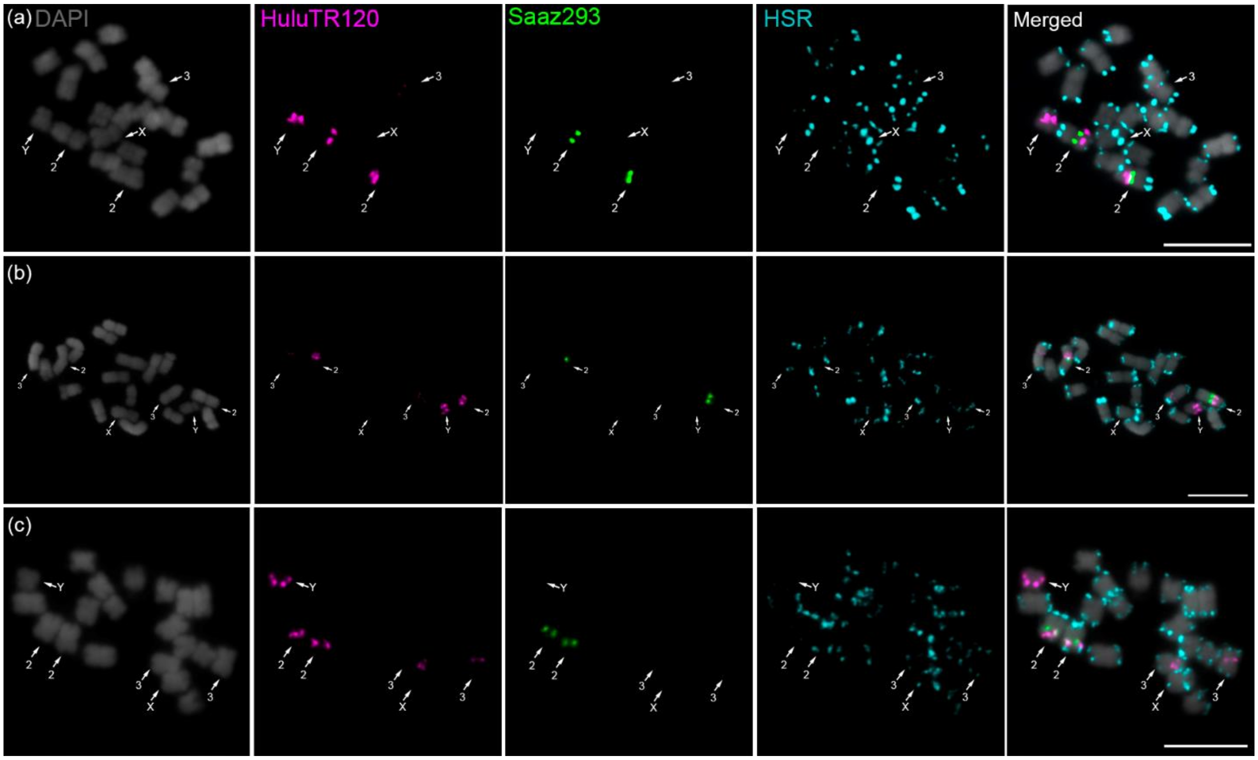

**Fig. S8** The localization of centromeric satellite Saaz40 (magenta) and 45S rDNA (green) on chromosome 6 in male (a) and female (b) of *Humulus lupulus*. Satellite Saaz40 exhibit the same pattern on metaphase chromosomes to that of satellite Saaz85 (Fig. 1g). Mitotic chromosomes were counterstained with DAPI. Scale bar = 10 μm.

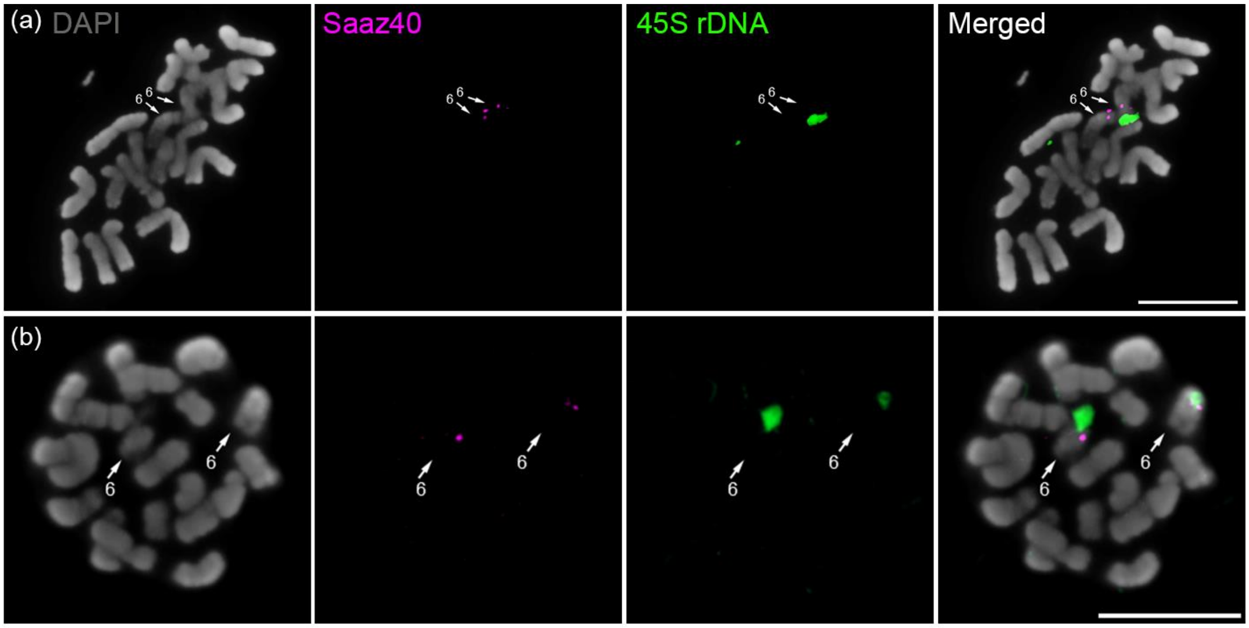

**Fig. S9** The localization of centromeric satellite Saaz85 (magenta) and 45S rDNA (green) on chromosome 6 in (a) male and (b) female *Humulus lupulus*. Subtelomeric repeat HSR (cyan) was used to distinguish X and Y chromosomes. Mitotic chromosomes were counterstained with DAPI. Scale bar = 10 μm.

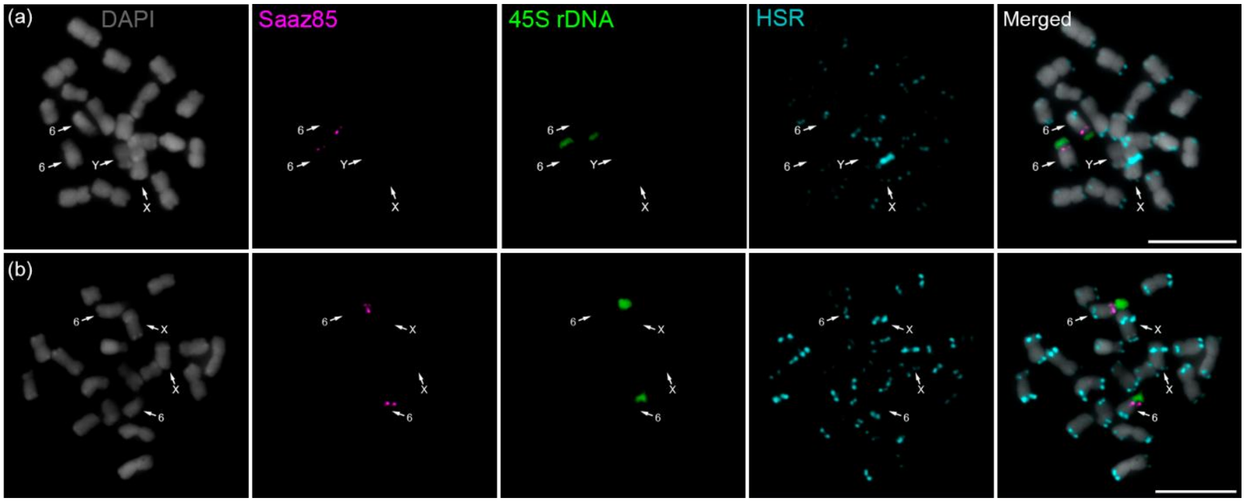

**Fig. S10** Chromosome pairing and organization during diakinesis of *Humulus lupulus* Lib male. (a) Subtelomeric satellite HSR (magenta) shows position of the PAR and locates pairing between X and Y chromosomes. (b) HuluTR120 (green) was used to differentiate Y chromosome. Meiotic chromosomes were counterstained with DAPI. Scale bar = 10 µm.

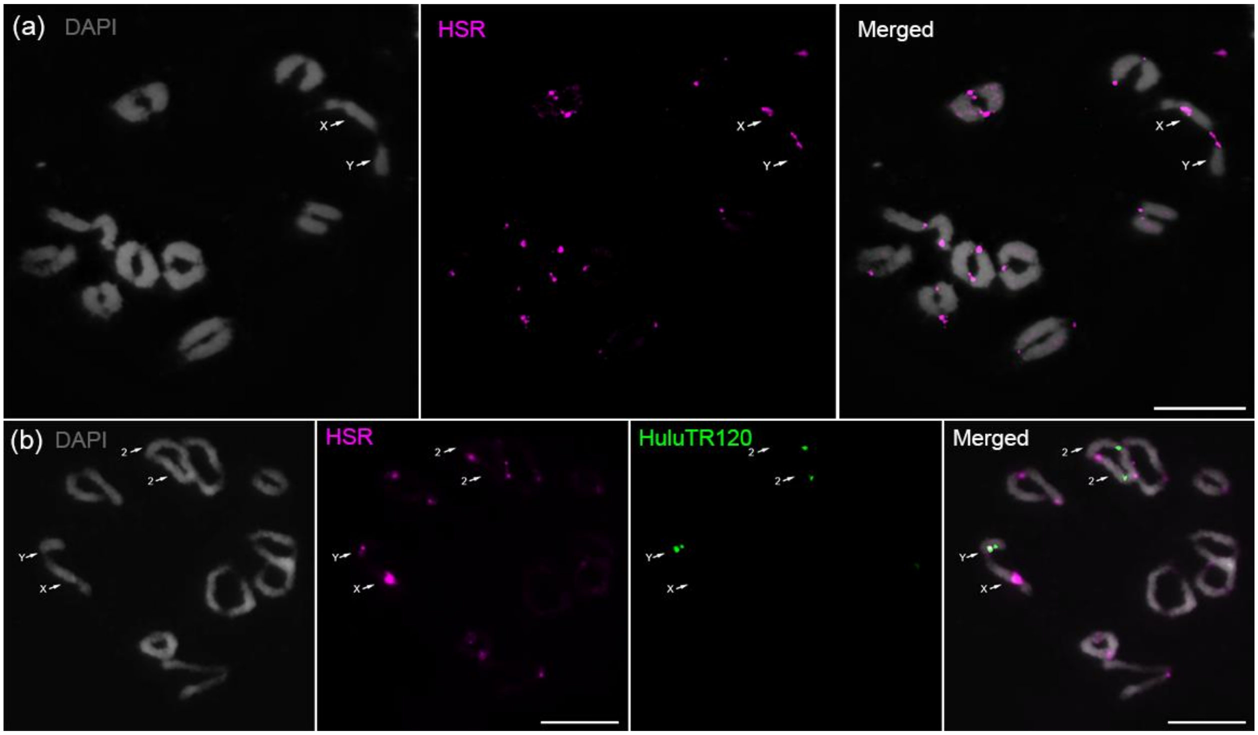

**Fig. S11** Distribution of major centromeric repeats on metaphase chromosomes of *Humulus lupulus* Saaz female. Subtelomeric repeat HSR (cyan) was used to distinguish two X chromosomes (pericentromeric localization on p-arm and subtelomeric localization on q-arm). (a) Localization of SaazCEN (magenta) and HSR (cyan). (b) Simultaneous localization of SaazCEN (magenta), SaazCRM1 (green), and HSR (cyan); (c) SaazCEN (magenta), Saaz293 (green), and HSR (cyan); (d) HuluTR120 (magenta), Saaz293 (green), and HSR (cyan); (e) SaazCEN (magenta), HuluTR120 (green), and HSR (cyan). Note the pericentromeric localization of HuluTR120 on chromosome pair 2 and chromosome 3 (only one chromosome from pair). (f) Localization of SaazCEN (magenta), satellite Saaz85 (green), and HSR (cyan). Chromosomes were counterstained with DAPI. Scale bar = 10 μm.

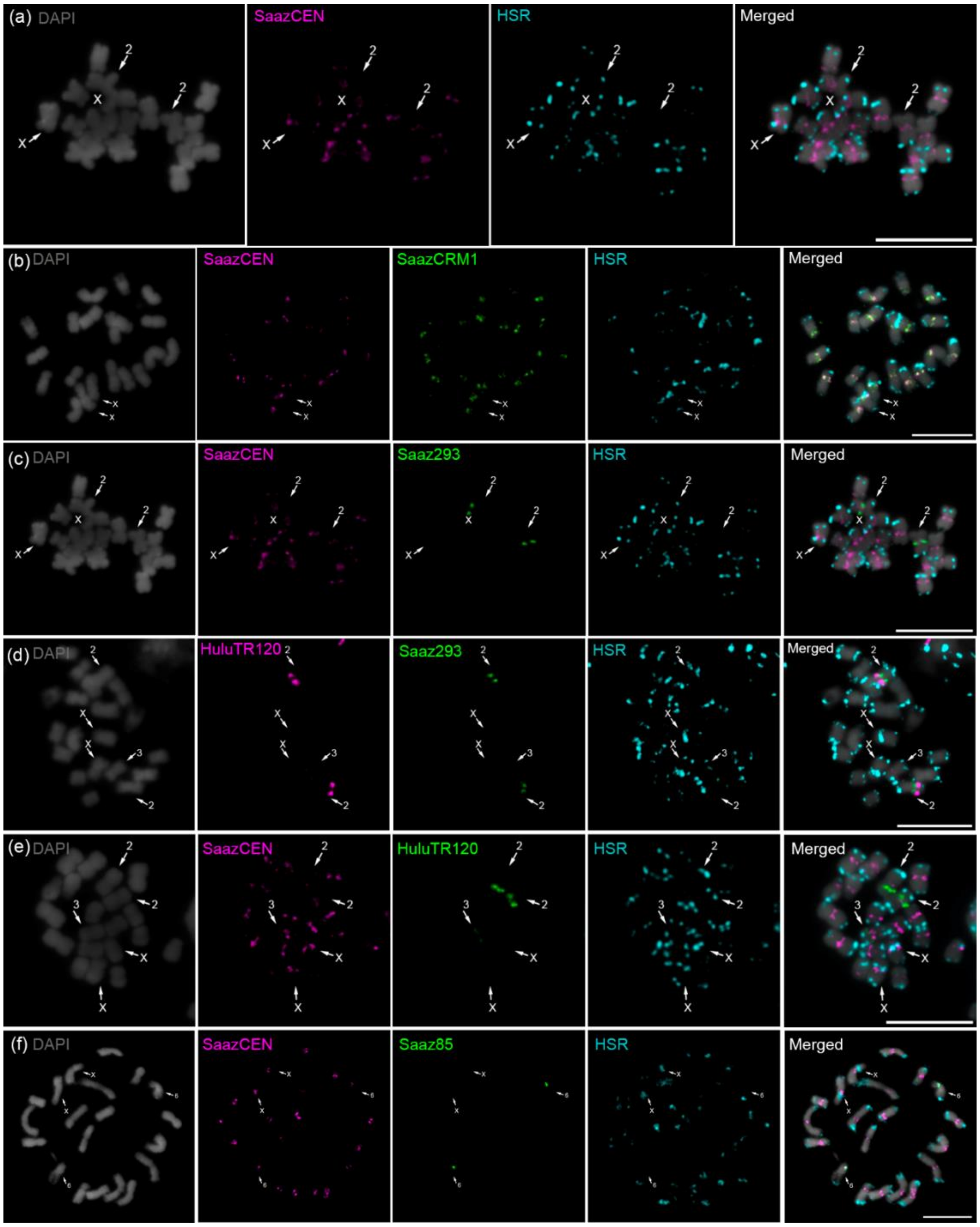

**Fig. S12** Distribution of major centromeric repeats on metaphase chromosomes of *Humulus lupulus* Lib male. Subtelomeric repeat HSR (cyan) was used to distinguish X (pericentromeric distribution on p-arm) and Y chromosomes (only p-arm). (a) Simultaneous localization of SaazCEN (magenta) and HSR (cyan); (b) SaazCEN (magenta), SaazCRM1 (green), and HSR (cyan); (c) SaazCEN (magenta), Saaz293 (green), and HSR (cyan); (d) HuluTR120 (magenta), Saaz293 (green), and HSR (cyan); (e) SaazCEN (magenta), HuluTR120 (green), and HSR (cyan); (f) SaazCEN (magenta), Saaz85 (green), and HSR (cyan). Chromosomes were counterstained with DAPI. Scale bar = 10 µm.

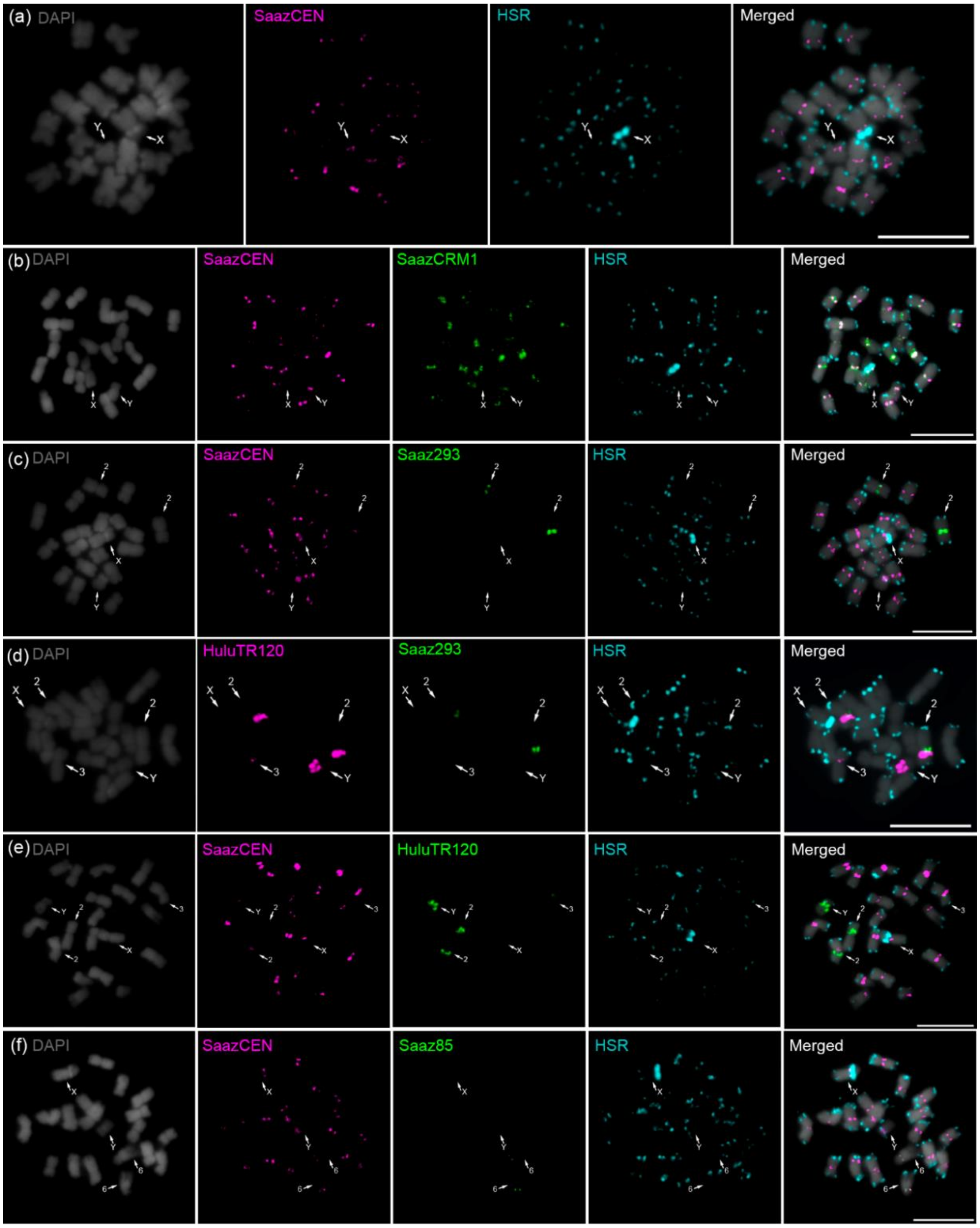

**Fig. S13** Dot plot and sequence similarity of three major centromeric satellites Saaz85, Saaz293 and Saaz40 of chromosomes 2, 3, 6, and 8. All three satellites show large conserved motifs within each sequence.

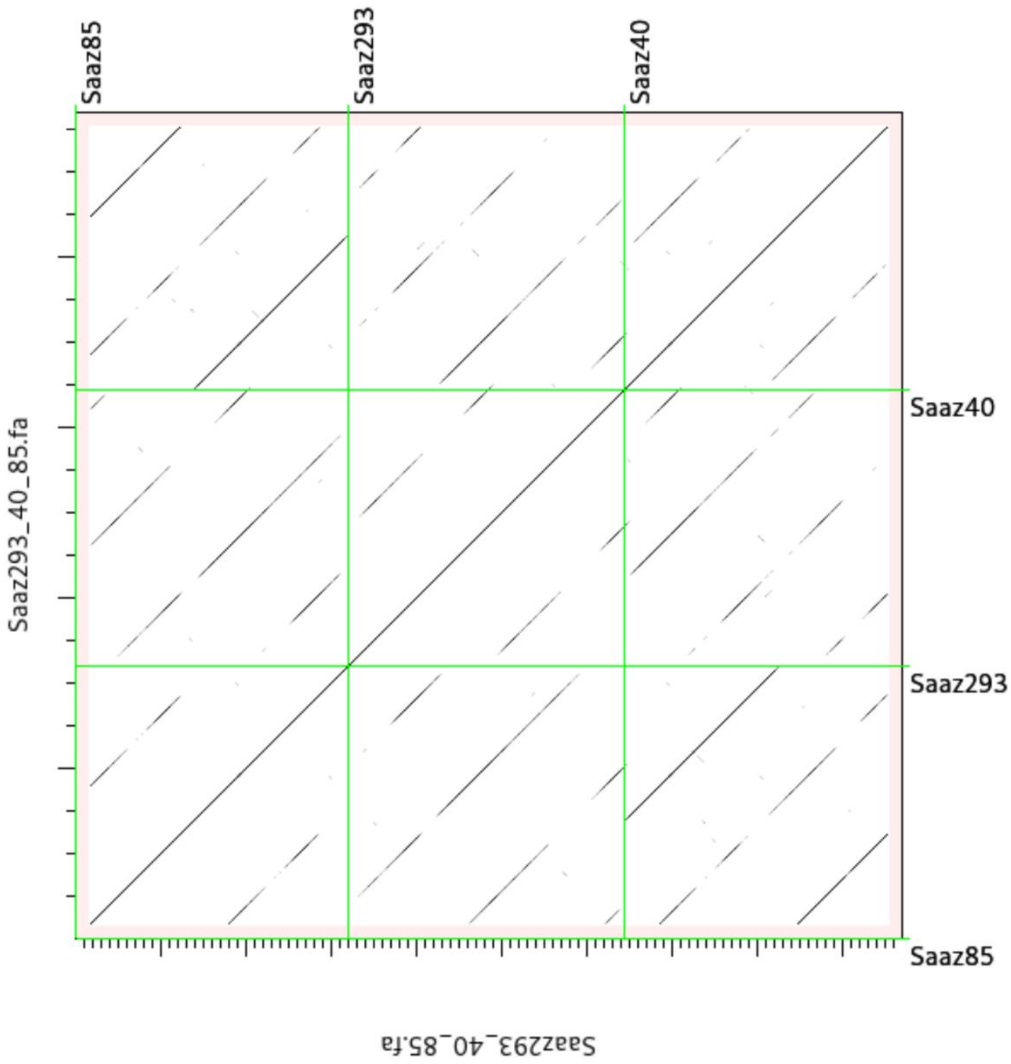

**Fig. S14** LTR retrotransposons composition of all centromeres of *Humulus lupulus* Saaz female. (a) Repeat composition of Ty1/*Copia* families. Ale (green) and Angela (orange) families are represented in centromeres all studied chromosomes and together consist main proportion of Ty1/*Copia* insertions. The other families (Ikeros, SIRE and TAR) are represented unequally. Family Ikeros (blue) is specific for centromeres of chromosomes 2, 3 and X. (b) Repeat composition of Ty3/*Gypsy* families. The most abundant families Tekay and CRM are represented in all chromosomes in similar proportions, except chromosome 3. Retand and Athila represent minor fraction on all chromosomes.

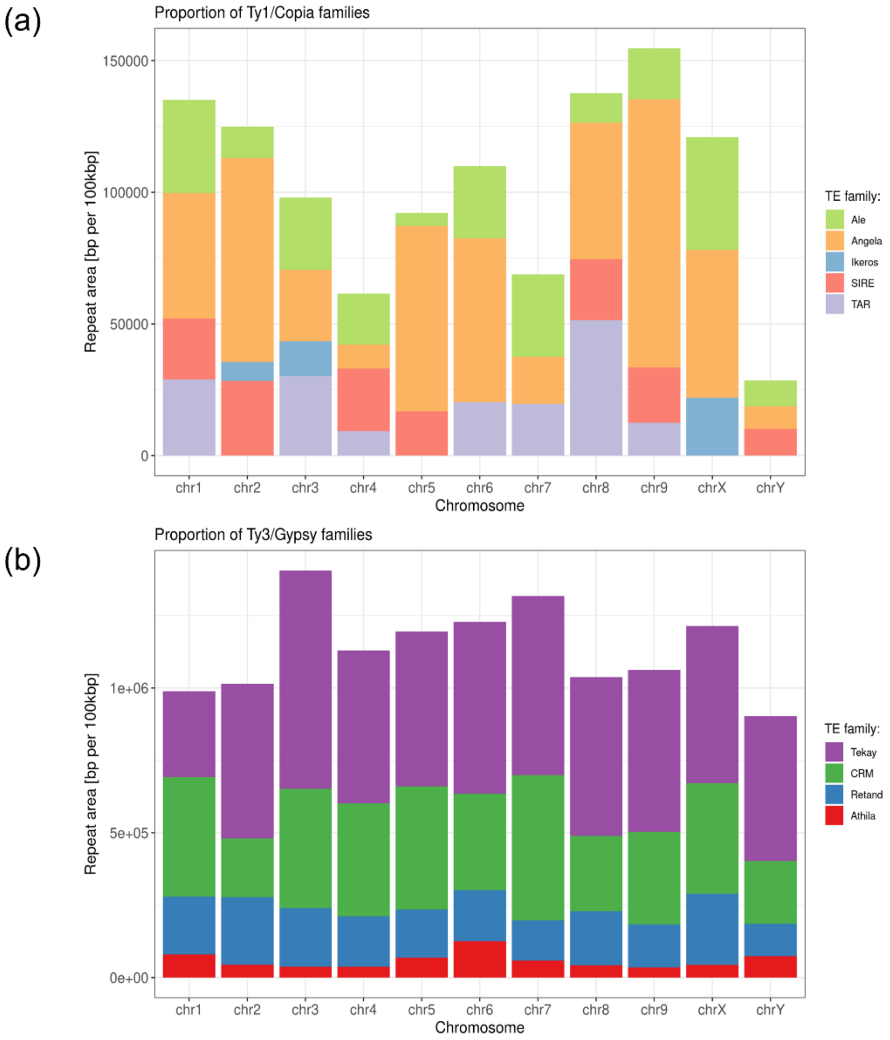

**Fig. S15** Distribution and estimated insertion time of Ty1/*Copia* and Ty3/*Gypsy* families of LTR retrotransposons for all chromosomes. The most recent CRM insertions within the HICENH3 binding domain. Note that chromosomes 1, 2, 3, 5, 7, and X possess large expanded HICENH3 positive region, with more recent CRM insertions within the main binding domain.

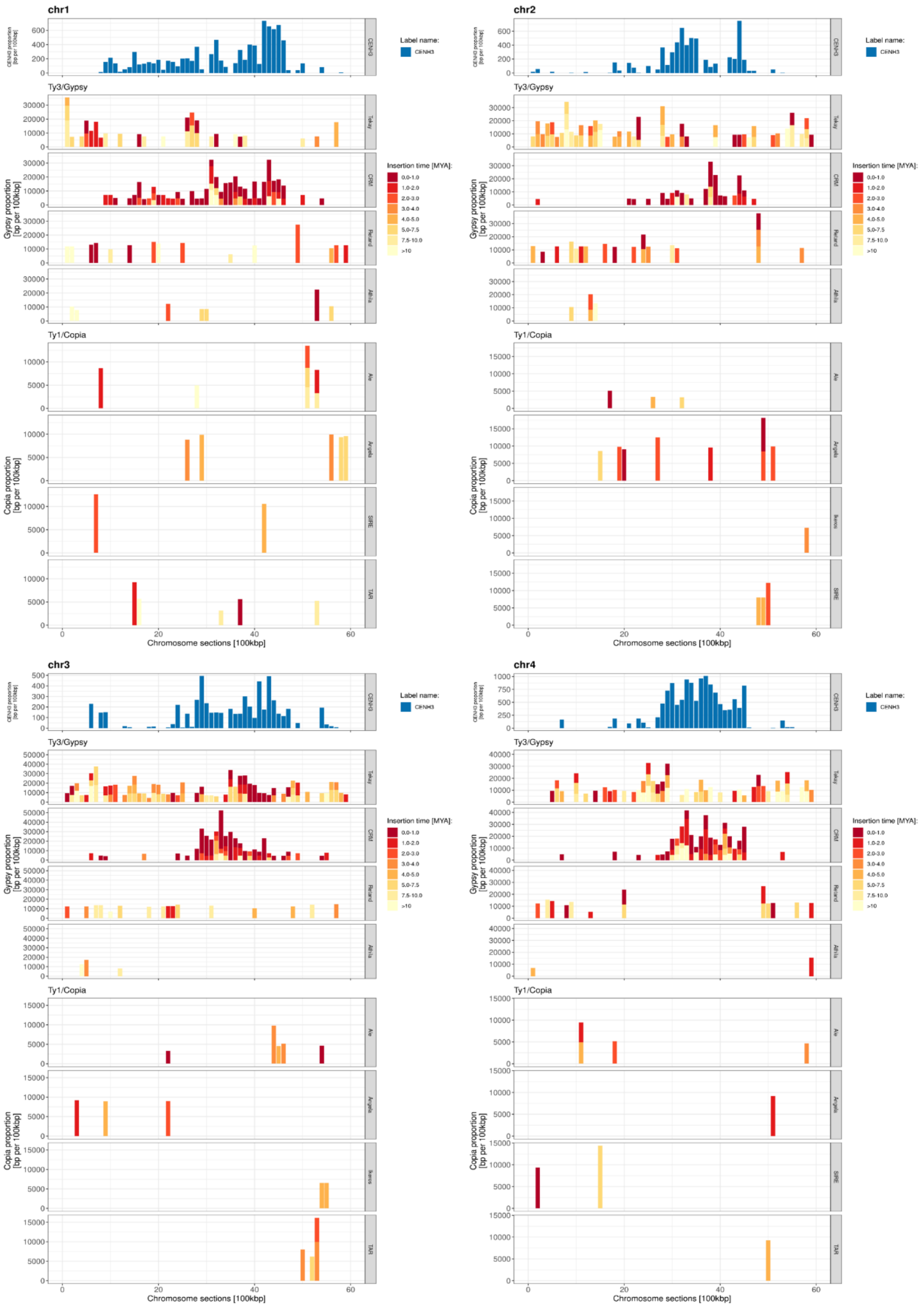

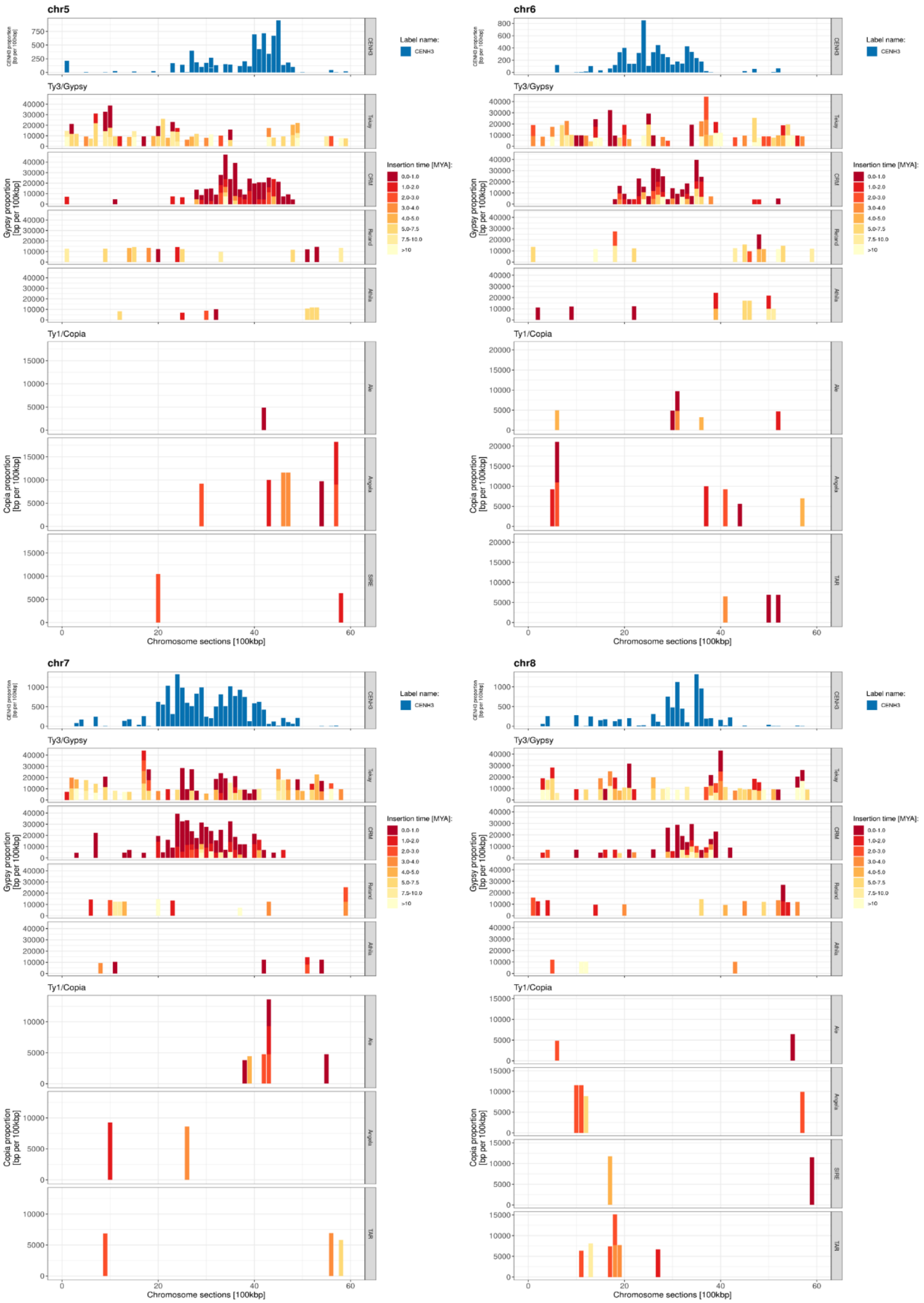

chr5
chr6
chr7
chr8
Label name:
CENH3
Ty3/Gypsy
Tekay
CRM
Reina
Athila
Insertion time [MYA]:
0.0-1.0
1.0-2.0
2.0-3.0
3.0-4.0
4.0-5.0
5.0-7.5
7.5-10.0
>10
Gypsy proportion [bp per 100kbp]
Ty1/Copia
Ale
Angela
SIRE
TAR
Copia proportion [bp per 100kbp]
Chromosome sections [100kbp]

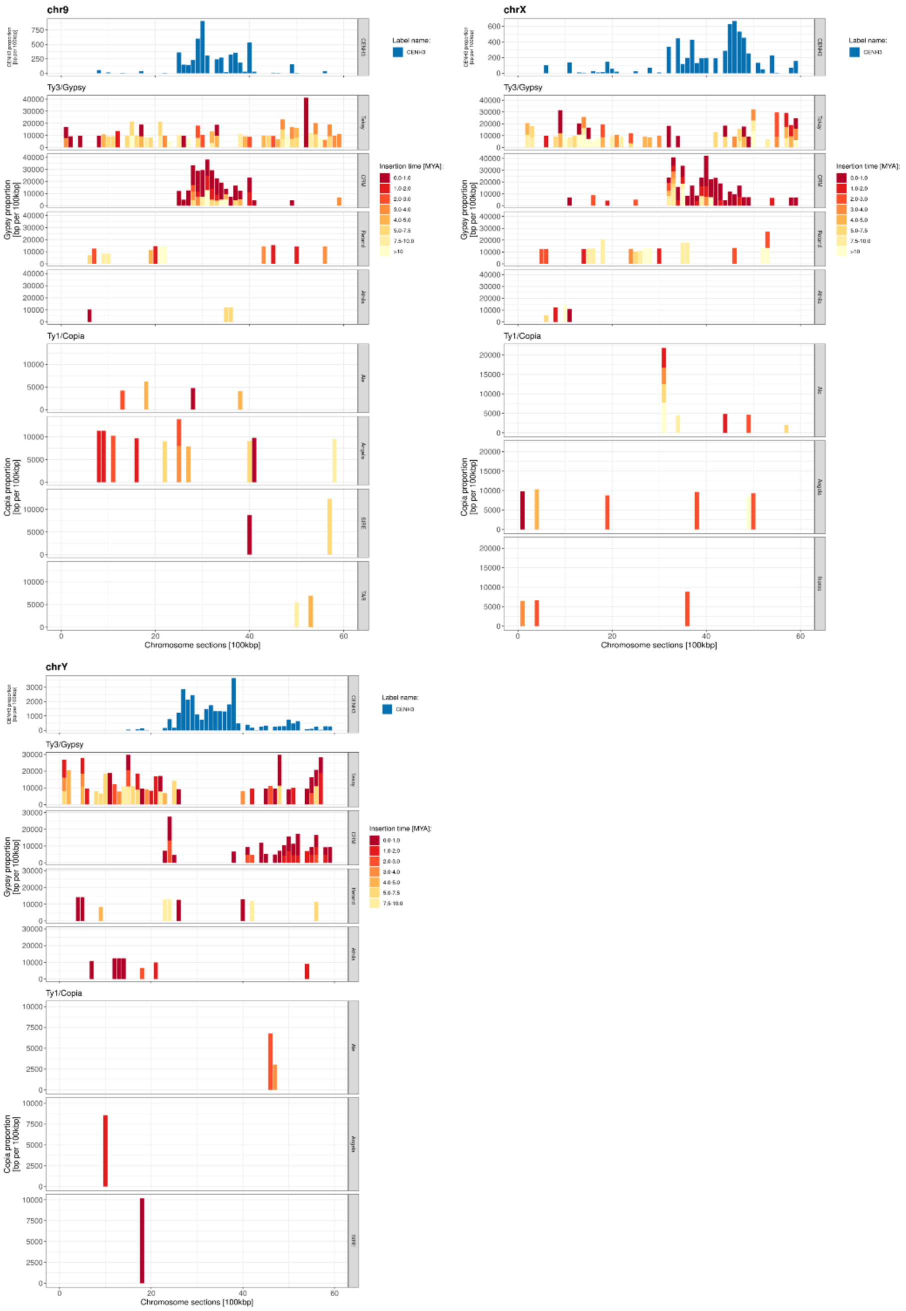
chr9
chrX
chrY
Ty3/Gypsy
Ty1/Copia
Label name:
CENH3
Insertion time [MYA]:
0.0-1.0
1.0-2.0
2.0-3.0
3.0-4.0
4.0-5.0
5.0-7.5
7.5-10.0
>10
Gypsy proportion [bp per 100kbp]
Copia proportion [bp per 100kbp]
Chromosome sections [100kbp]

**Fig. S16** The insertion time of autonomous (red), dominant non-autonomous (green), and minor non-autonomous CRMs (blue). Chromosomes 4 and X have the oldest insertions of both, autonomous and non-autonomous (dominant and minor) CRM groups.

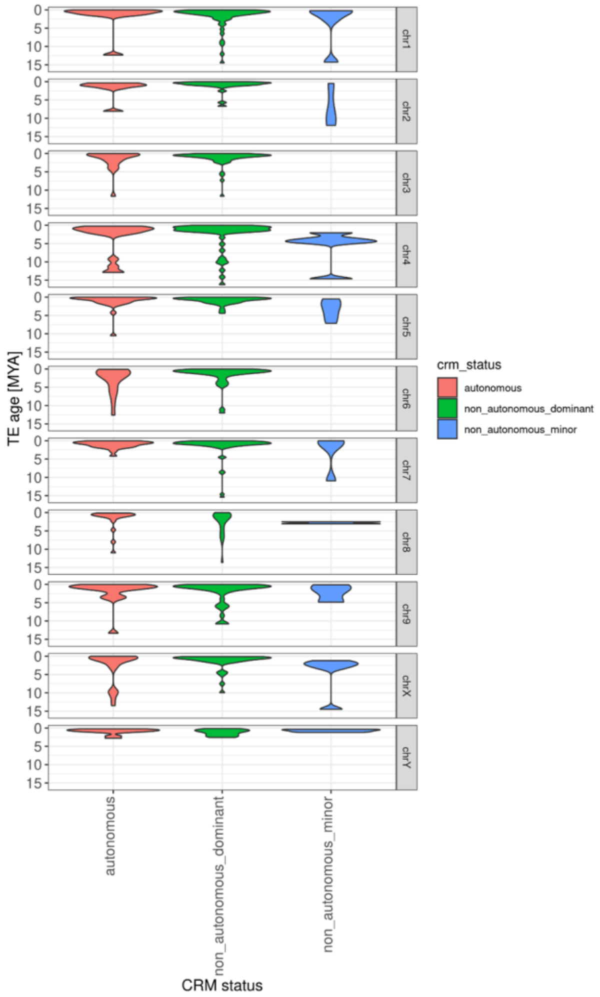

**Fig. S17** The full-length distribution of centromeric Ty3/*Gypsy* CRM retrotransposons in *Humulus lupulus.* Autonomous CRMs (red) with complete sets of domains (Fig. S16) exhibited longer lengths than both groups of nonautonomous CRMs – dominant (green) and minor (blue).

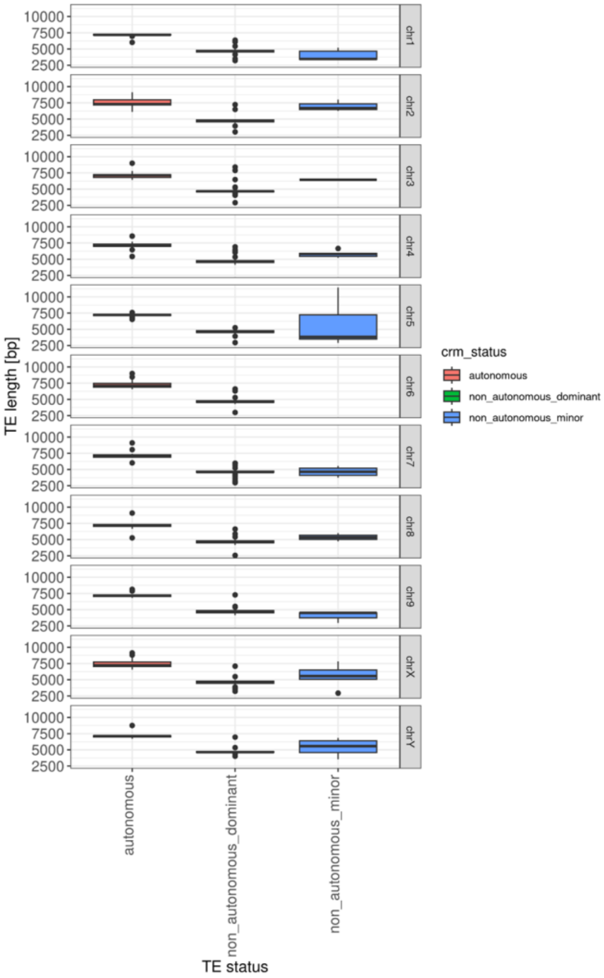

**Fig. S18** (a) Phylogenetic analysis of the autonomous and non-autonomous CRM retrotransposons correlated with insertion age (MYA). (b) The most abundant CRM lineages with recent insertions were found in non-autonomous elements (labeled by green oval).

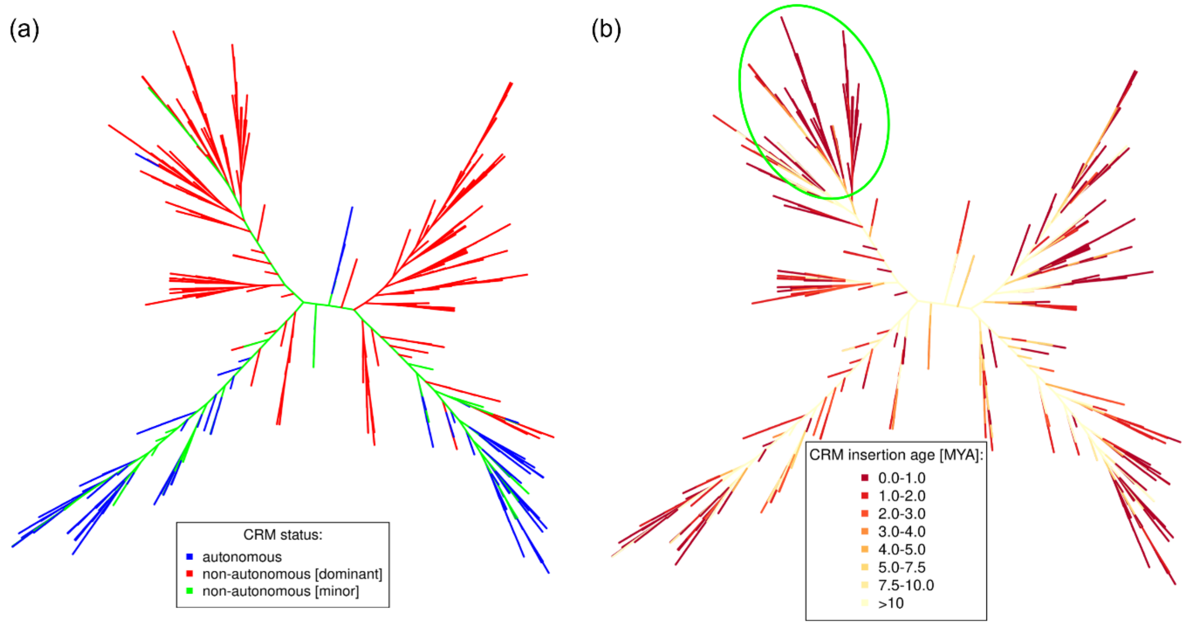

**Fig. S19** Distribution of SaazCEN and SaazCRM1 within the autonomous and non-autonomous CRM retrotransposons in *Humulus* centromere. The majority of CRMs (94.4%) contain both SaazCEN and SaazCRM1 repeats (purple). A minority of CRMs (3.6%) contain only SaazCRM1 (orange). Note longer distribution of both types of repeats within the centromeric region of chromosome 1 and HICENH3 binding region (Fig. S14).

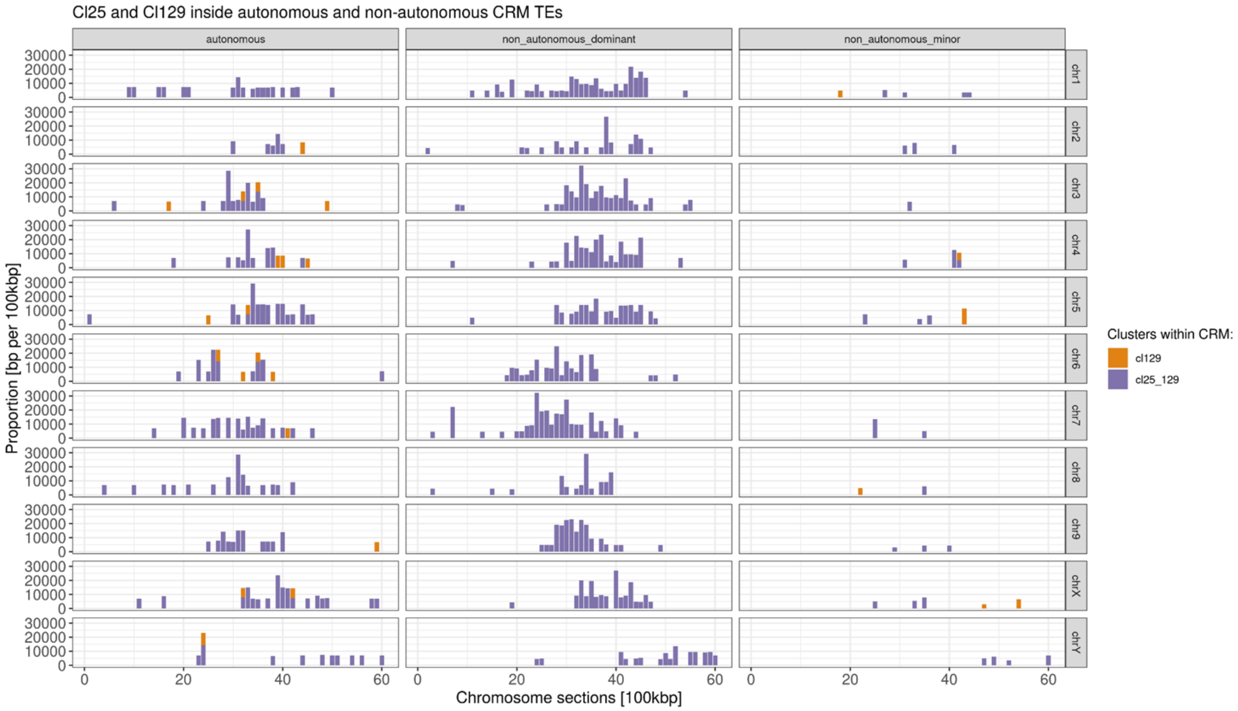

**Fig. S20** Localization of HlCENH3 within CRM retrotransposons in *Humulus* centromere. The most binding regions of CRM retrotransposons are found within spacer and LTRs. Other domains show low or no binding of HlCENH3.

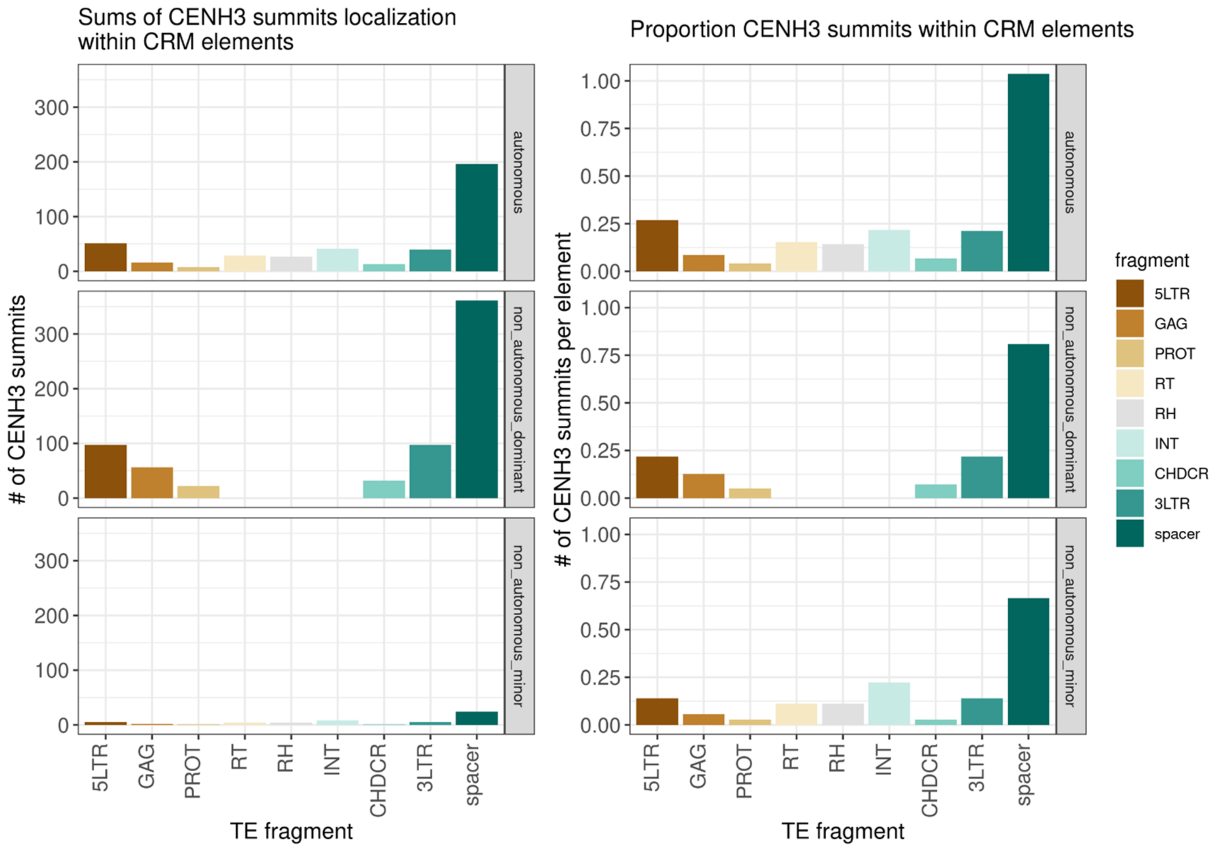

**Fig. S21** Localization of the major centromeric satellites on chromosomes 2 and 6. Chromosome 2 is clearly differentiated by the localization of HuluTR120 (red), 5S rDNA (green), and Saaz293 (cyan) repeats. Chromosome 6 is differentiated based on the localization of HSR satellite (magenta) and 45S rDNA (yellow). Dash line shows the position of 45S rDNA on the second chromosome 6. Mitotic chromosomes were counterstained with DAPI. Scale bar = 10 μm.

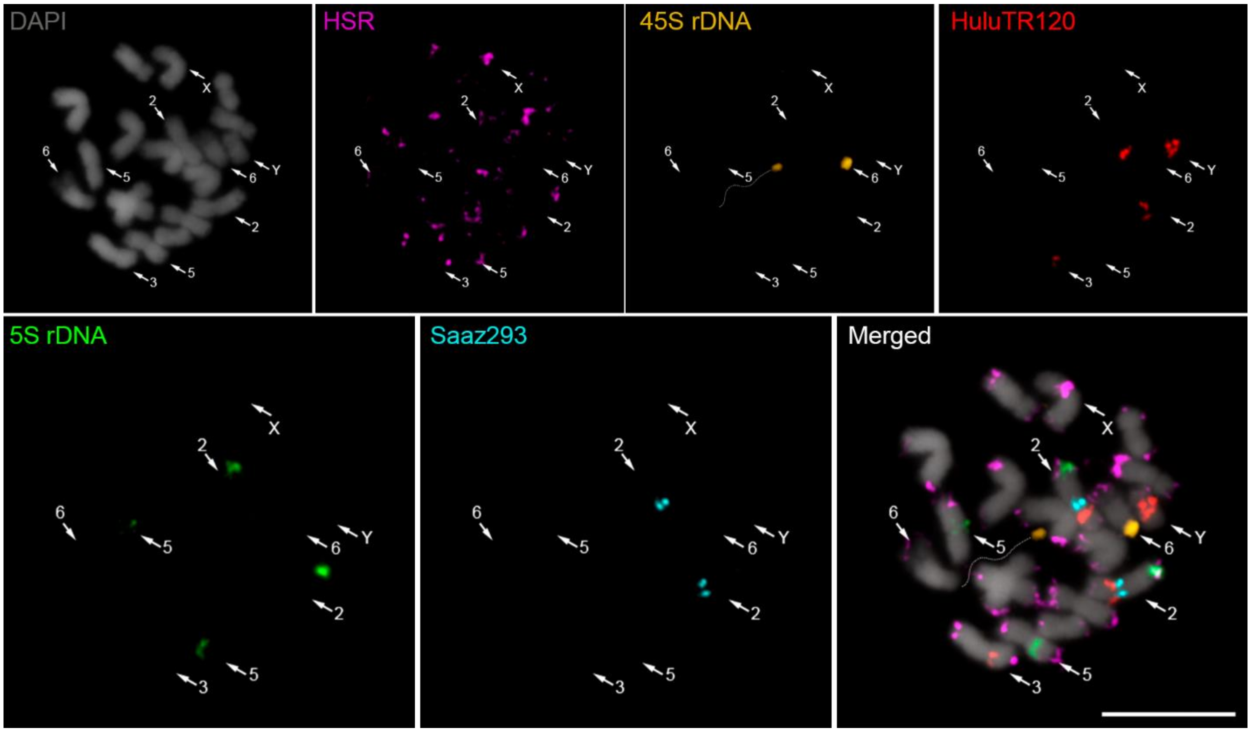

**Fig. S22** Positioning of chromosome 2 within an interphase nucleus of male and female *Humulus lupulus*. (a) Two distinct loci of satellite Saaz293 in the diploid (2n = 20) male and (b) female nucleus of *Humulus lupulus*. Arrows indicate Saaz293 signal on chromosome 2 (green). The number of HuluTR120 signals (magenta) corresponds to number of signals on metaphase chromosomes in diploid cells, on chromosomes 2, 3 and Y in Fig. 1d. Note the peripheral localization of both satellite clusters. Nuclei were counterstained with DAPI. Scale bar = 10 μm.

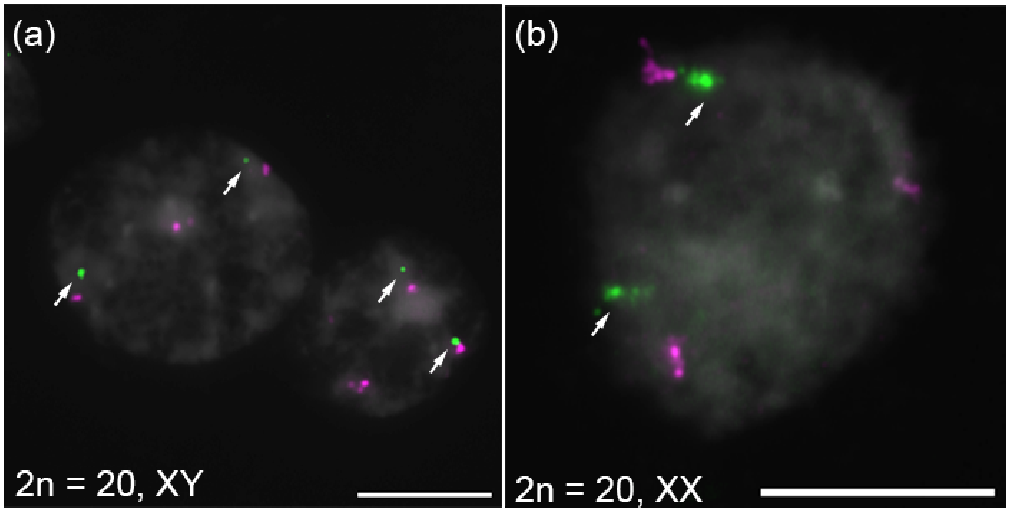

**Table. S1** List of *Humulus lupulus* plants utilized in this study.

| Hop cultivars | Chromosome number | Collected material |
|---|---|---|
| Saaz hop - Osvald's clone 72 | 2n = 20, XX | Leaves |
| Liběšice male (Lib male) | 2n = 20, XY | Leaves, panicles |
| 15246 | 2n = 20, XY | Leaves |
| 15249 | 2n = 20, XY | Leaves |
| 15276 | 2n = 20, XY | Leaves |
| $F_1$ progeny (Osvald's clone 72 x Liběšice male) | 2n = 20, XY | Panicles |
| Wild hop – Horní Heršpice | 2n = 20, XY | Panicles |

**Table S2** List of primers used for PCR and preparation of FISH probes.

| Primer name | Forward primer (5´-3´) | Reverse primer (5´- 3´) | GenBank Accession | References |
|---|---|---|---|---|
| HlCENH3 | CCTCCTCTTCCTCCACTCCA | TTTCCCTCCAAGTCGACGTG | | |
| Saaz293 | ACAAATTCAATGGTGGCCGT | TCCCTGATATCTTTCTTTCTCCCT | | |
| SaazCEN | GGGTTGTCTTTAAATTTCGGT | CAACCCTTTGTCTTCCTAATGA | | |
| SaazCRM1 | GGGCATGTGAGAGAAAGCATG | TACAACAAAGCCTAGGAAAACAAGC | | |
| Saaz85 | CCTTGTTCGGGATTTATTGAATCA | GTGGGTTAAACGAGTGAAAAAGA | | |
| Saaz40 | GGTCCGAGGTAGTGAGTTGTG | TCAAAATTTGTGCAGAAATCGGT | | |
| HuluTR120 | AGTTCCTGGATATAACCAGGTC | GAAAACTTAGTTCGTGTTAACT | MN537570 | (Easterling et al. 2020) |
| HSR1 | CCCTCTGGTGAATTGGAGAT | CCTTTCAGAAATCTTCGATTTCTCTA | GU831574 | (Divashuk et al. 2011) |
| 5S rDNA | GTTTTCAGGGGTGCAACACG | CTTACGGCTCAAAAGTTTGT | MN537579 | (Easterling et al. 2020) |
| 45S rDNA | TGCCCGTTGCTCTGATGATT | TCCACCAACTAAGAACGGCC | AF223066.1 | |

**Table S3** Enzyme mixture used for digestion of young leaves.

| Enzyme mixture | |
|---|---|
| 1% pectolyase | P3026, Sigma-Aldrich |
| 0.75% cellulase R-10 Onozuka | C8001.0005, DuchefaBiochemie |
| 0.75% cellulase | 219466, Sigma-Aldrich |
| 1% cytohelicase | C8274, Sigma-Aldrich |

**Table S4** Clusters of tandem repeats selected for FISH including ChipSeq Mapper outputs.

| Repeats | Monomer Length (bp) | ChIP Hits | Input Hits | Normalized ratio ChIP/Input | Annotation | Reference |
|---|---|---|---|---|---|---|
| **SaazCEN** | 284 | 89401 | 6006 | 31.0 | Satellite | |
| **Saaz293** | 323 | 206894 | 6777 | 63.6 | Satellite | |
| **Saaz85** | 178 | 22376 | 137676 | 0.34 | Satellite | |
| **Saaz40** | 324 | 42832 | 906 | 98.5 | Satellite | |
| **HuluTR120** | 120 | 402834 | 36562 | 23.0 | Satellite | MN537570 (Easterling et al. 2020) |
| **SaazCRM1** | | 713546 | 37509 | 39.6 | Ty3/Gypsy _CRM | |

**Table S5** Genomic fraction of repetitive DNA in the *H. lupulus* Saaz and Lib male genome estimated from genomic abundance of reconstructed contigs by RepeatExplorer2 pipeline.

| **Repeats** | **Superfamily** | **Lineage** | **Clade** | **Genome proportion (%) in *H. lupus*** | | |
|---|---|---|---|---|---|---|
| | | | | **Female** | **Male** | **Average** |
| LTR retrotransposons | Ty1/Copia | Ale | | 0.10 | 0.09 | 0.10 |
| | | Alesia | | 0.02 | 0.02 | 0.02 |
| | | Angela | | 7.98 | 7.90 | 7.94 |
| | | Bianca | | 0.01 | 0.00 | 0.01 |
| | | Ikeros | | 0.35 | 0.33 | 0.34 |
| | | Ivana | | 0.01 | 0.02 | 0.02 |
| | | SIRE | | 0.82 | 0.76 | 0.79 |
| | | TAR | | 2.44 | 2.35 | 2.40 |
| | | Tork | | 0.04 | 0.04 | 0.04 |
| | Total Ty1/Copia | | | **11.77** | **11.51** | **11.64** |
| | Ty3/Gypsy | Chromovirus | CRM | 0.20 | 0.21 | 0.21 |
| | | | Galadriel | 0.21 | 0.20 | 0.21 |
| | | | Tekay | 21.60 | 21.60 | 21.60 |
| | | Non-chromovirus | Athila | 2.53 | 2.54 | 2.54 |
| | | | Retand | 15.13 | 15.26 | 15.20 |
| | Total Ty3/Gypsy | | | **39.67** | **39.81** | **39.74** |
| Total LTR retrotransposons | | | | **51.44** | **51.32** | **51.38** |
| DNA transposons | | EnSpm_CACTA | | 2.49 | 2.34 | 2.42 |
| | | MuDR_Mutator | | 0.43 | 0.42 | 0.43 |
| | | PIF_Harbinger | | 0.04 | 0.05 | 0.05 |
| | | hAT | | 0.05 | 0.04 | 0.05 |
| | Total DNA transposons | | | **3.01** | **2.85** | **2.93** |
| Tandem repeats | Satellite | | | 0.35 | 0.32 | 0.34 |
| | rDNA | | | 0.52 | 0.60 | 0.56 |
| Organelle | | | | 2.36 | 3.42 | 2.89 |
| Unclassified | | | | 6.51 | 6.29 | 6.40 |
| **Total** | | | | **64.19** | **64.80** | **64.50** |

**Table S6** Tandem repeats in *H. lupulus* Saaz and Lib male genome identified by RepeatExplorer2 pipeline.

| **Cluster** | **Supercluster** | **Monomer Length (bp)** | **HL MF ratio** | **Annotation** | **Accession** | **Reference** |
|---|---|---|---|---|---|---|
| HSR1 (Cl124) | 60 | 383 | 0.89 | Satellite | GU831574 | Divashuk *et al.*, 2011 |
| HSR0 (Cl304) | 211 | 178 | 1.23 | Satellite | MH188533.1 | Easterling *et al.*, 2018 |
| Saaz293 (Cl293) | 200 | 323 | 1.32 | Satellite | | |

**Table S7** Number and frequency of meiotic abnormalities in three male accessions of *Humulus lupulus*.

| **Meiotic abnormalities** | **Lib male** | **F1 progeny (Osvald's clone 72 x Lib male)** | **Wild hop** |
|---|---|---|---|
| **Regular tetrad stage** | 1260 (87.50%) | 1530 (87.28%) | 1852 (89.43%) |
| **Unreduced cells** | 133 (9.24%) | 94 (5.36%) | 129 (6.23%) |
| **Unviable microspore** | 36 (2.50%) | 122 (6.96%) | 72 (3.48%) |
| **Micronuclei** | 11 (0.76%) | 7 (0.40%) | 18 (0.87%) |
| Total | **1440** | **1753** | **2071** |

**Tab. S8** Number and frequency of viable and unviable pollen grain in three male accessions of *Humulus lupulus*.

| **Pollen viability** | **Lib male** | **F1 progeny (Osvald's clone 72 x Lib male)** | **Wild hop** |
|---|---|---|---|
| **Viable** | 4909 (96.05%) | 3994 (99.45%) | 4413 (99.9%) |
| **Unviable** | 202 (3.95%) | 22 (0.55%) | 4 (0.09%) |
| Total | **5111** | **4016** | **4417** |
